# Prothrombin Knockdown Protects Podocytes and Reduces Proteinuria in Glomerular Disease

**DOI:** 10.1101/2023.06.20.544360

**Authors:** Amanda P. Waller, Katelyn J. Wolfgang, Iva Pruner, Zachary S. Stevenson, Eman Abdelghani, Kaushik Muralidharan, Tasha K. Wilkie, Angela R. Blissett, Edward P. Calomeni, Tatyana A. Vetter, Sergey V. Brodsky, William E. Smoyer, Marvin T. Nieman, Bryce A. Kerlin

## Abstract

Chronic kidney disease (CKD) is a leading cause of death, and its progression is driven by glomerular podocyte injury and loss, manifesting as proteinuria. Proteinuria includes urinary loss of coagulation zymogens, cofactors, and inhibitors. Importantly, both CKD and proteinuria significantly increase the risk of thromboembolic disease. Prior studies demonstrated that anticoagulants reduced proteinuria in rats and that thrombin injured cultured podocytes. Herein we aimed to directly determine the influence of circulating prothrombin on glomerular pathobiology. We hypothesized that (pro)thrombin drives podocytopathy, podocytopenia, and proteinuria. Glomerular proteinuria was induced with puromycin aminonucleoside (PAN) in Wistar rats. Circulating prothrombin was either knocked down using a rat-specific antisense oligonucleotide or elevated by serial intravenous infusions of prothrombin protein, which are previously established methods to model hypo- (LoPT) and hyper-prothrombinemia (HiPT), respectively. After 10 days (peak proteinuria in this model) plasma prothrombin levels were determined, kidneys were examined for (pro)thrombin co-localization to podocytes, histology, and electron microscopy. Podocytopathy and podocytopenia were determined and proteinuria, and plasma albumin were measured. LoPT significantly reduced prothrombin colocalization to podocytes, podocytopathy, and proteinuria with improved plasma albumin. In contrast, HiPT significantly increased podocytopathy and proteinuria. Podocytopenia was significantly reduced in LoPT vs. HiPT rats. In summary, prothrombin knockdown ameliorated PAN-induced glomerular disease whereas hyper-prothrombinemia exacerbated disease. Thus, (pro)thrombin antagonism may be a viable strategy to simultaneously provide thromboprophylaxis and prevent podocytopathy-mediated CKD progression.

## INTRODUCTION

Kidney disease is the 8^th^ leading cause of death in the U.S. and cardiovascular disease is the leading cause of mortality in patients with chronic kidney disease (CKD) (*1–9*). Glomerular diseases are a leading cause of CKD and end-stage kidney disease (ESKD) (*6, 10*). The glomerular filtration barrier is composed of three layers: (1) fenestrated endothelial cells, (2) collagen basement membrane, and (3) podocytes which interdigitate to form the slit diaphragm (a specialized adherence junction) (*11*). Proteinuria, glomerular podocyte injury (podocytopathy) leading to podocyte loss (podocytopenia), and glomerulosclerosis are hallmarks of glomerular disease-mediated CKD (*1, 4, 12–16*). Podocytes are terminally differentiated epithelial cells that maintain glomerular endothelial cell phenotype via paracrine signaling, synthesize basement membrane components, and mechanically resist pulsatile intracapillary blood pressure (*11, 17, 18*). Podocyte function is thus critical to glomerular filtration barrier maintenance and podocyte injury and/or death leads to urinary loss of plasma proteins (proteinuria) (*18–21*). While proteinuria due to acute podocyte dysfunction is potentially reversible, it becomes irreversible with ≥20% podocyte depletion and glomerulosclerotic lesions form on the extraluminal surface of glomerular capillaries in the areas of podocyte loss (*1, 12, 22*). Proteinuria, in turn, is a key driver of CKD progression, leading to renal tubular inflammation and interstitial fibrosis (*1*). Thus, podocyte preservation and proteinuria constraint are crucial to delay or prevent both CKD progression and CKD-related cardiovascular disease.

Nephrotic syndrome (NS) is comprised of a group of primary glomerular diseases that collectively are a leading cause of ESKD, the most severe form of CKD (*6*). NS is clinically characterized by massive proteinuria, hypoalbuminemia, edema, and dyslipidemia (*23, 24*). The monogenic, familial forms of NS are predominantly due to mutations in podocyte-specific genes and idiopathic NS is typified by podocytopathy (*1, 25*). Rodent models of podocytopathy-mediated proteinuria have thus become important tools for the investigation of NS pathogenesis and CKD progression (*10, 12*). The massive proteinuria of NS includes urinary loss of albumin and other plasma proteins, including coagulation system zymogens, cofactors, and inhibitors with variable synthetic compensation that manifests as an acquired hypercoagulopathy (*26, 27*). Thus, NS is itself a profoundly prothrombotic condition with increased risk for both venous and arterial thrombosis, even before CKD onset (*8, 26, 28–33*).

We and others have recently shown that prothrombin (*F2*) may directly injure podocytes during NS (*34–36*). *In vitro* experiments demonstrated that thrombin injured conditionally immortalized rat podocytes in a protease-activated receptor-dependent manner (*34*). Moreover, inhibition of thrombin with natural anticoagulant proteins or pharmacologic anticoagulants reduced proteinuria in two mechanistically rat NS models: puromycin aminonucleoside (PAN) and transgenic podocyte-specific human diphtheria toxin receptor rats (*34, 36–38*). Moreover, prothrombin colocalization to podocytes *in vivo* was proportional to proteinuria, strongly suggesting that prothrombin, originating from the plasma compartment, interacts with podocytes to drive podocytopathy, podocytopenia, and proteinuria during NS. What remains unknown is if thrombin injures *in vivo* podocytes during proteinuria or if anticoagulants exert their antiproteinuric effects by an alternative mechanism. The objective of this study was thus to test the hypothesis that thrombin drives *in vivo* podocyte injury using the well-established PAN-NS model (*10, 34, 36, 39*). Here we demonstrate that plasma prothrombin levels modulate both *in vivo* podocyte health and proteinuria during PAN-NS. These data suggest that anticoagulant thrombin inhibition may simultaneously reduce the risk of thrombotic cardiovascular disease and preserve kidney function during NS (*40*).

## MATERIALS AND METHODS

### Animals

All experimental rat protocols were approved by the Institutional Animal Care and Use Committee at the Abigail Wexner Research Institute, in accordance with the NIH Guide for the Care and Use of Laboratory Animals. Wistar rats (body weight ∼150 g, age ∼45-50 d) were used for all experiments (**Table S1**). Transgenic rat *F2* hypo- and hyper-expressing models are not yet available (*41–44*). However, antisense oligonucleotides (ASOs) have been used to successfully knock-down mouse prothrombin expression to levels comparable to those seen in *F2*^lox/-^ knockdown mice (*45–48*). Thus, ASOs provide an alternative means to achieve *in vivo* prothrombin knockdown. Intravenous administration of human prothrombin protein has been used to model hyper-prothrombinemia in mice (*49*). We thus utilized ASO-mediated prothrombin knockdown and serial prothrombin infusions to mimic hypo- and hyper-prothrombinemia (LoPT and HiPT), respectively. Subcutaneous ASO or intravenous prothrombin was administered at the indicated doses and frequencies to achieve LoPT and HiPT conditions, respectively (**Figure S1)**. Proteinuria was induced with a single tail vein injection of PAN (MilliporeSigma, Burlington, MA; 50 mg/kg diluted in saline) on Day 0 (*10, 34, 39*). Anti-*F2* ASO (401027) treatment (60 mg/kg subcutaneous twice weekly) was begun on Day −17.5 to induce LoPT prior to onset of podocyte injury and continued through Day +9. Prothrombin infusions were begun on Day 0, immediately following PAN (or saline) administration. Control rats received twice weekly scrambled ASO (141923) on Days - 17.5 to +9, saline tail vein injections on Days 0, 3, 6, and 9 (instead of PAN and/or prothrombin). Sham rats were treated identically to control rats but did receive PAN on Day 0. Morning spot urine samples were collected on Days 0 (before PAN or saline; **Figure S2**) and 10 to determine urinary protein-to-creatinine ratio. Following Day 10 urine collection, the rats were anesthetized with 3% isoflurane and blood was collected from the inferior vena cava through a 23-G needle into final concentration 0.32% NaCitrate/1.45 µM Corn Trypsin Inhibitor (CTI; Prolytix, Essex Junction, VT) and processed to Platelet Poor Plasma (PPP), as previously described (*39*). Urine and plasma were stored at −80°C until analyzed. Following exsanguination, both kidneys were removed, decapsulated, and placed in ice cold PBS. One quarter kidney was set aside for immunohistology and glomeruli were isolated from the remainder of both kidneys.

### Glomerular Isolation and Podocyte Flow Cytometry

Glomeruli were isolated from renal cortex using a standard sequential sieving method, washed and resuspended in ice cold PBS, then manually counted in a hemocytometer, as previously described (*50–52*). An aliquot of the glomerular suspension (∼2,500 glomeruli) was dissociated to a single cell suspension in Digestion Buffer at 37°C on a 1400/min shaker with intermittent shearing through an 18-G needle for 26 minutes. Tissue debris was removed with a 40-micron filter, the cells were washed with 10 mL HBSS, centrifuged at 1500 rpm for 5 min, then resuspended in 1 mL HBSS supplemented with 0.1% BSA and 4’,6-diamidino-2-phenylindole (DAPI; 1 μg/mL). The cells were then prepared for flow cytometry analysis by staining with Live/Dead near IR dye and FITC-conjugated synaptopodin antibody (Fitzgerald, North Acton, MA; 1:5) in Perm/Wash buffer. The cells were analyzed on a LSR II flow cytometer (BD Biosciences) and synaptopodin expression was gated only on cells with a DAPI-positive intact nucleus. Isotype controls, unstained cells, and an open channel were used to identify and calibrate for autofluoresence. Synaptopodin-positive podocytes were normalized to bead standards to estimate podocyte counts which were then divided by the starting number of glomeruli to enumerate podocytes/glomerulus. Reassuringly, we observed a direct linear relationship between podocyte counts determined by flow cytometry and a histologic method (see Supplement: Materials and Methods, **Figure S3**, and **Videos 1-4**) (*53*). A second aliquot of the glomerular cell suspension was stained with FITC-TUNEL (1:5) and APC-conjugated synaptopodin antibody (1:4) to determine % TUNEL-positive podocytes. Flow cytometry data points for each individual rat represents an analysis of 100,000 events.

### Immunofluorescence Colocalization

Paraffin-embedded rat kidney sections were deparaffinized and processed for synaptopodin and thrombin co-immunofluorescence, as previously described (*34*). However, the sections were photobleached for 72 hours prior to deparaffinization using a custom-built lightbox to reduce background autofluorescence (*54*). Lightbox construction was adapted from previous descriptions using mirrored acrylic panels and a broad-spectrum LED panel (*55–57*). Images were captured with a Leica DMI 4000B inverted fluorescence microscope (Deerfield, IL) and analyzed with ZEN Black software (Zeiss USA, Thornwood, NY). Colocalization analysis was performed on a pixel-by-pixel basis, wherein every pixel within the defined region of interest is plotted in a scattergram based on fluorescence intensity of each channel. The software calculates the colocalization coefficient as the proportion of colocalized pixels (positive on both channels) within the total population of a single channel, green (synaptopodin-positive) for the purposes of this study (*58*).

### Coagulation Parameters

Plasma prothrombin concentration was measured with rat- and human-specific ELISA (MyBioSource Inc, San Diego, California). Prothrombin activity was determined chromogenically using a commercially available assay (Rox Prothrombin; DiaPharma, West Chester, OH), as previously described (*59*), and are expressed as a percentage of rat pooled normal plasma (rPNP). Endogenous thrombin potential (ETP) was determined on a 1:1 dilution of PPP with the Technothrombin TGA kit (Technoclone, Vienna, Austria) using TGA RC low reagent, and read on a Spectramax M2 fluorescent plate reader (Molecular Devices, Sunnyvale, California), as previously described (*39, 59*).

### Statistical Analyses

One- or two-way ANOVA (analysis of variance) for multiple group comparisons, using SigmaStat software (Systat, San Jose, CA). When a significant difference was identified by ANOVA, post hoc tests were performed using the Student-Newman-Keuls technique. Statistical significance was defined as *P<*0.05. Figures were prepared using GraphPad Prism (Boston, MA). Data are presented as mean ± SE.

**See Supplement for additional Materials and Methods details.**

## RESULTS

### Antisense Oligonucleotide-Mediated Hypo-prothrombinemia in Healthy Rats

ASO 401027, which is 100% complementary to a 3’ segment of rat *F2* mRNA, effectively abolished *F2* mRNA expression in primary rat hepatocytes vs. control ASO (0.00±0.00 vs. 1.0±0.01 relative fold change; *P*<0.001; **Figure S4**), whereas ASO 401025 (75% complementary to rat *F2* mRNA) less effectively suppressed *F2* expression (0.63±0.14 relative fold change; *P*=0.024). ASO 401027 also reduced prothrombin activity in primary rat hepatocyte protein lysates to 1.41±0.16 vs. 9.13±0.13 mIU/mL in control ASO lysates (*P*<0.001), whereas ASO 401025 less efficiently reduced activity to 2.31±0.10 mIU/mL (*P*<0.001). In dose finding studies, ASO 401027 was administered subcutaneously to healthy rats twice weekly for 17.5 days (**Figure 1)**. These data suggested that maximal plasma prothrombin activity reduction (∼32% of rPNP) was achieved with 60 mg/kg/dose. To mimic the planned full-length experiment, this dose (60 mg/kg twice weekly) was then continued for 27.5 days after which plasma prothrombin activity was reduced to 31.6±3.52% (vs. 100.3±1.45% in rPNP; *P*<0.001) and endogenous thrombin potential (ETP) was reduced to 514.3±58.7 nM*min (vs. 2451.0±140.5 nM*min; *P*<0.001). Thus, whereas prothrombin activity was reduced by ∼68%, ETP was reduced by ∼71% suggesting that circulating prothrombin reduction impaired thrombin-dependent coagulation cascade amplification (*60*). Hepatic synthesis is the major source of circulating prothrombin and hepatic *F2* mRNA expression was essentially abolished by this 27.5 day ASO treatment regimen (*P*<0.01) (*61, 62*). Thus, ASO 401027 effectively induced sustained LoPT in otherwise healthy rats.

**Figure 1:**
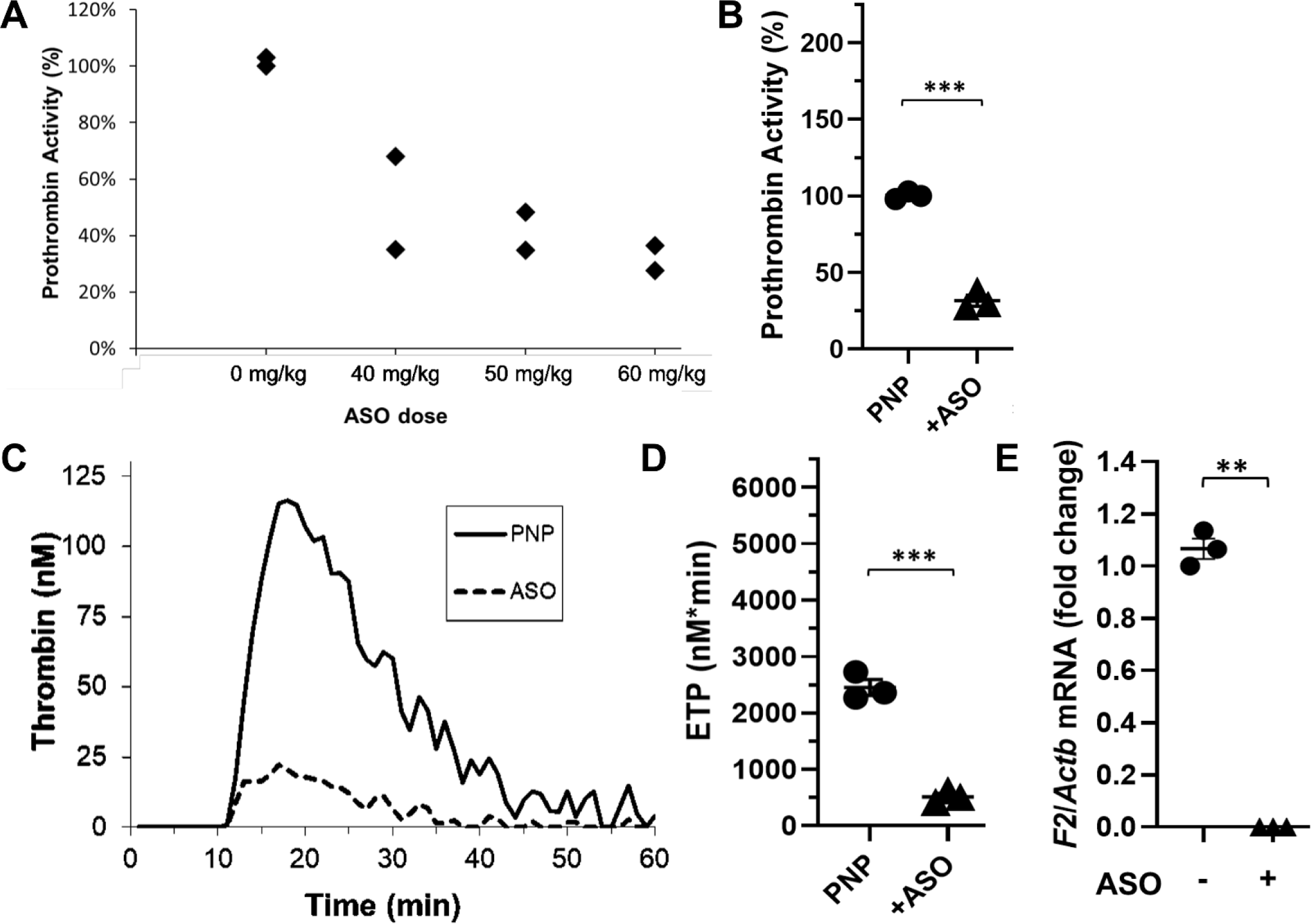
Antisense Oligonucleotide-Mediated Hypo-prothrombinemia in Healthy Rats. (**A**)Twice weekly treatment with ASO 401027 for 17.5 days reduced plasma prothrombin enzymatic activity in a dose-dependent manner that appeared to plateau at ∼60 mg/kg/dose (relative to rat pooled normal plasma prothrombin activity; *n*=2 per dose level). Twice weekly ASO 401027 at 60 mg/kg/dose significantly reduced plasma prothrombin enzymatic activity (**B**) and endogenous thrombin potential (ETP, **C**, **D**) measured at day 27.5 (*n*=3). (**E**) RT-qPCR demonstrated that twice weekly ASO 401027 at 60 mg/kg/dose significantly reduced day 27.5 hepatic prothrombin (*F2*) gene expression (relative to β-actin (*Actb*)) (*n*=3). \*\*\**P*<0.001

### Serial Prothrombin Infusion-Mediated Hyper-prothrombinemia in Healthy Rats

The hemostatic system is highly conserved amongst vertebrate species, including mammals (*63, 64*). Thus, rPNP thrombin generation was augmented by addition of human prothrombin as expected (**Figure S5**). Moreover, rat thrombomodulin effectively regulated the spiked-in human prothrombin. These observations are similar to those previously observed using human prothrombin added to mouse PNP and indicate that human prothrombin is compatible with rat pro- and anti-coagulant systems (*49*). In preliminary experiments, intravenously administered human prothrombin (31.25 mg/kg) generated peak plasma prothrombin activity of ∼207% 1-hour post-dose with an apparent half-life of ∼61 hours (**Figure 2**), which is similar to that previously reported in humans (*65*). To mimic the planned full-length experiment, 31.25 mg/kg prothrombin was administered as a loading dose on day 0 with maintenance doses of 16.67 mg/kg on days 3, 6, and 9. On day 10, 24 hours after the day 9 dose, prothrombin activity was 186.8±10.4% (vs. 100.3±1.45% in rPNP; *P*<0.001). Similarly, day 10 ETP was elevated to 5766.0±207.0 (vs. 2451.0±140.5 nM*min; *P*<0.001). Thus, whereas prothrombin activity was enhanced by ∼86%, ETP was enhanced by ∼135% suggesting that elevated circulating prothrombin enhanced thrombin-dependent coagulation cascade amplification. These data demonstrate that human prothrombin induces hyper-prothrombinemia (HiPT) in otherwise healthy rats for up to 10 days.

**Figure 2:**
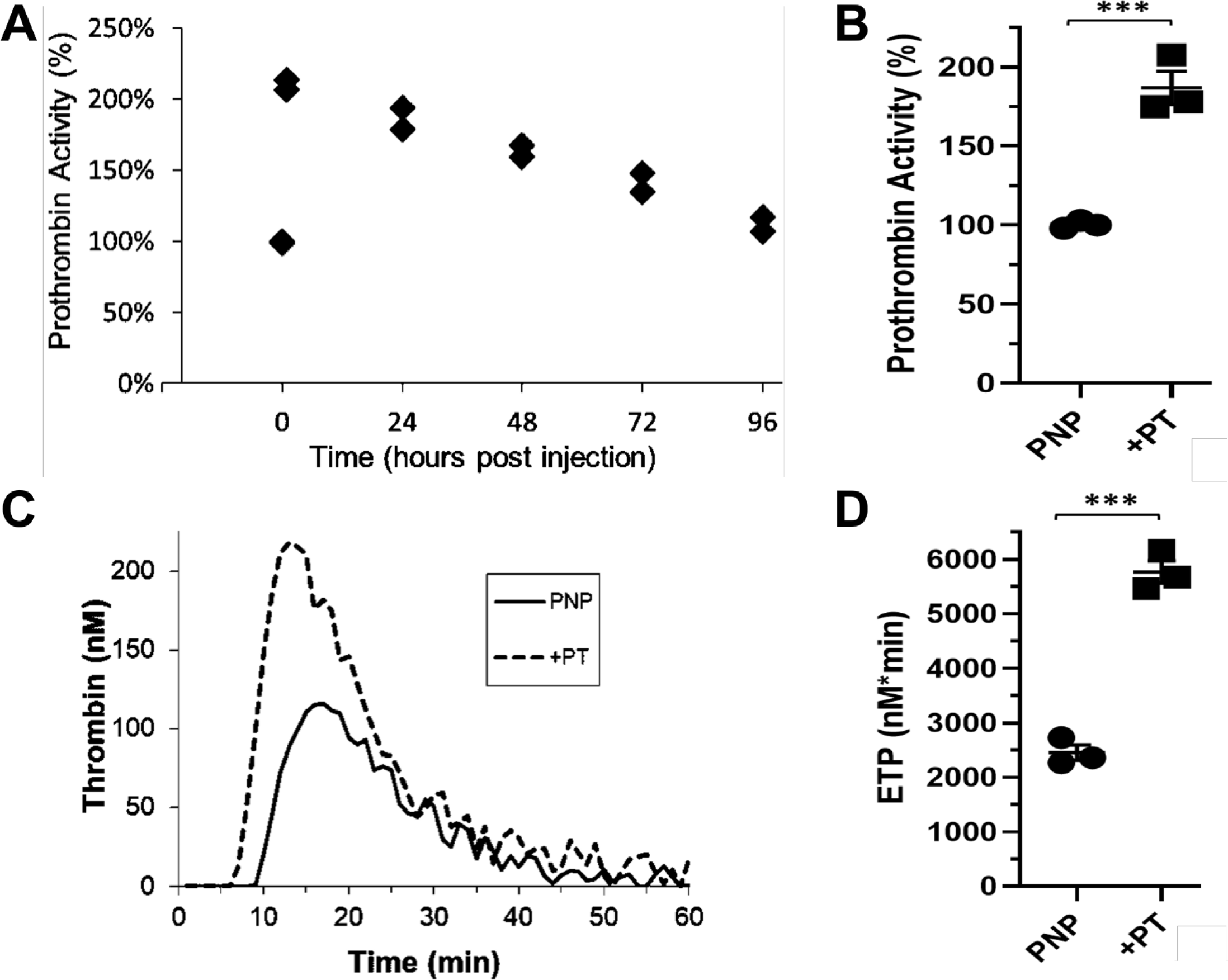
Serial Prothrombin Infusion-Mediated Hyper-prothrombinemia in Healthy Rats. (**A**) A single dose of intravenous prothrombin (31.25 mg/kg) increased plasma prothrombin enzymatic activity to a peak of ∼207% at 1-hour post-dose with an apparent half-life of ∼61 hours (relative to rat pooled normal plasma prothrombin activity; *n*=2 per time point). This 31.25 mg/kg loading dose was then followed by maintenance doses of 16.67 mg/kg on days 3, 6, and 9 resulting in significantly increased plasma prothrombin enzymatic activity (**B**) and endogenous thrombin potential (ETP, **C**, **D**) on day 10 (*n*=3). \*\*\**P*<0.001

### Prothrombin Modulation during Puromycin Aminonucleoside-Induced Rat Nephrotic Syndrome

Peak NS disease activity occurs at day 8-11 following PAN administration, we therefore evaluated prothrombin levels at day 10 to ensure that the manipulations observed above in otherwise healthy rats were sustained during PAN-mediated NS (PAN-NS) (*10, 34, 39*). Similar to the primary rat hepatocyte and *in vivo* dose finding studies described above, ASO 401027 treated rats (LoPT) had significantly reduced hepatic *F2* expression (*P*<0.05 vs Control) whereas *F2* expression was not significantly affected by PAN (Sham; *P*=0.35 vs Control) or prothrombin infusions (HiPT; *P*=0.28 and *P*=0.98 vs. Control and Sham, respectively; **Figure 3**). PAN-NS rats treated with ASO 401027 (LoPT) exhibited significantly reduced day 10 plasma prothrombin concentrations (0.92±0.10 μM) vs. Control (4.07±0.17 μM) and Sham (4.20±0.15 μM) whereas prothrombin infusions (HiPT) resulted in significantly elevated concentrations (8.16±0.38 μM; **Figures S6 and 3**). The LoPT and HiPT changes in prothrombin concentrations translated into significant changes in chromogenic prothrombin activity: LoPT 29.5±3.9% and HiPT 222.3±12.8% vs. Control (105.3±1.2%) and Sham (123.6±7.7%; **Figure 3**). Congruently, ETP was reduced to 306.7±62.9 nM*min in LoPT (vs. 2893.0±122.0 and 3417.0±235.8 nM*min in Control and Sham rats respectively; *P*<0.001) whereas HiPT rats had significantly elevated ETP (4406.0±407.2 nM*min; *P*<0.05). These data demonstrate successful manipulation of plasma prothrombin levels during rat PAN-NS.

**Figure 3:**
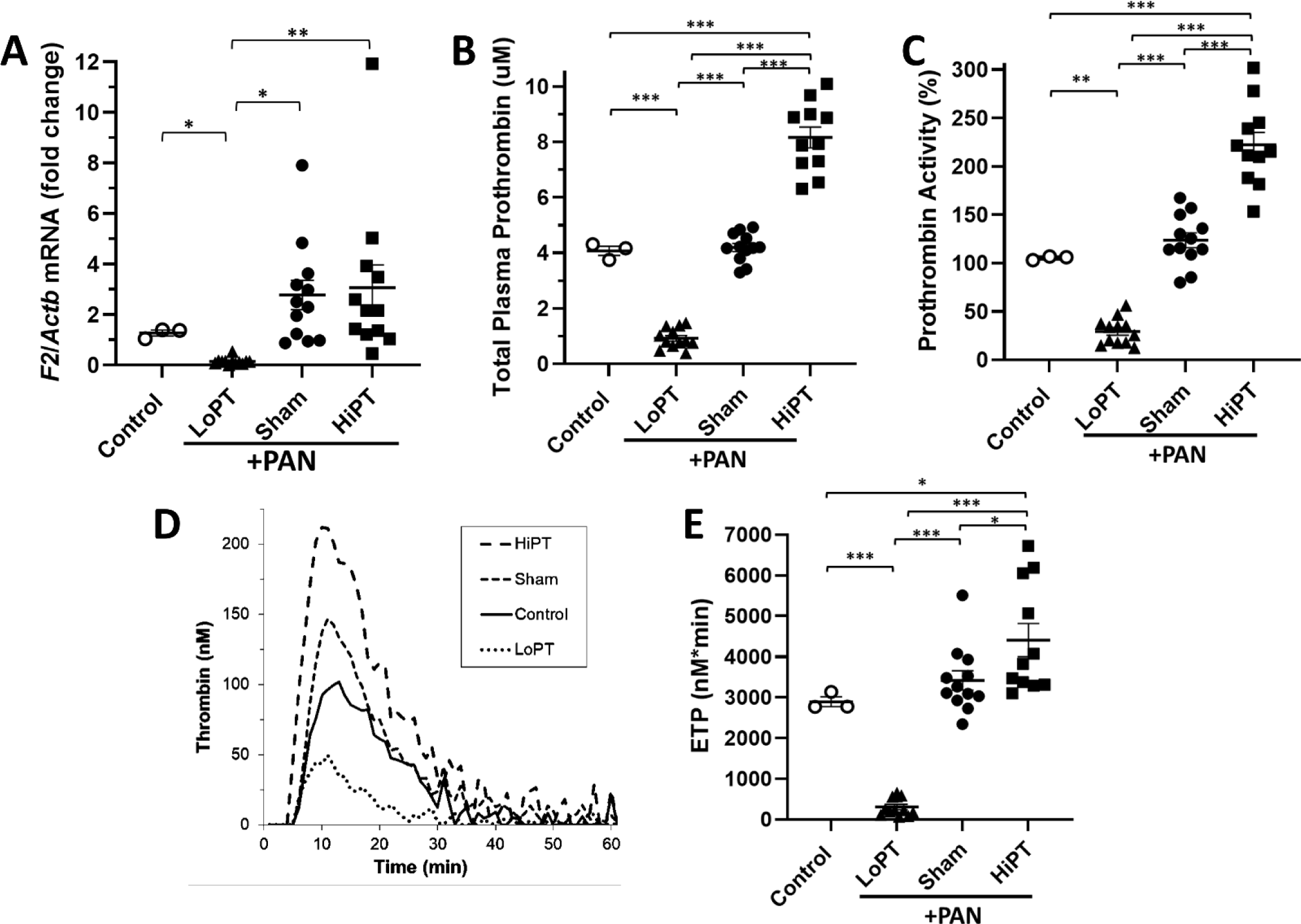
Prothrombin Modulation during Rat PAN-NS. On day 10, (**A**) RT-qPCR revealed that ASO 401027 significantly reduced hepatic prothrombin (*F2*) transcript levels (relative to β-actin (*Actb*)) whereas *F2* expression was not significantly altered by PAN-induced proteinuria or prothrombin infusions (*n*=3-12). Plasma prothrombin protein quantity (**B**), enzymatic activity (**C**), and endogenous thrombin potential (ETP, **D**, **E**) were significantly reduced in LoPT rats and significantly increased in HiPT rats in comparison to both control rats and PAN-NS (Sham) rats. *n*=3-12; \**P*<0.05, \*\**P*<0.01, \*\*\**P*<0.001; ○: Control; ▴: LoPT (ASO-mediated hypo-prothrombinemia); ●: Sham; ▪: HiPT (prothrombin infusion-mediated hyper-prothrombinemia)

### Plasma Prothrombin Levels Dictated Thrombin-Podocyte Interactions during Puromycin Aminonucleoside-Induced Rat Nephrotic Syndrome

We previously demonstrated that thrombin colocalized to podocytes during rat PAN-NS (*34*). We thus predicted that thrombin-podocyte interactions would fluctuate with plasma prothrombin levels. As expected, thrombin-podocyte colocalization was significantly reduced in LoPT rats (7.8±1.2% vs. 18.9±1.8% in Sham; *P*<0.01) to levels similar to Control (5.2±1.5%; *P*=0.30; **Figure 4**). Meanwhile, thrombin-podocyte colocalization was significantly higher than Control in both Sham (*P*<0.01) and HiPT (18.1±3.4%; *P*<0.05) rat PAN-NS. The latter data suggest that thrombin interactions with podocyte-expressed protease-activated receptors may be saturated at physiologic prothrombin concentrations (*34*). Nonetheless, these data strongly suggest that plasma prothrombin levels dictate podocyte exposure to thrombin during glomerular proteinuria.

**Figure 4:**
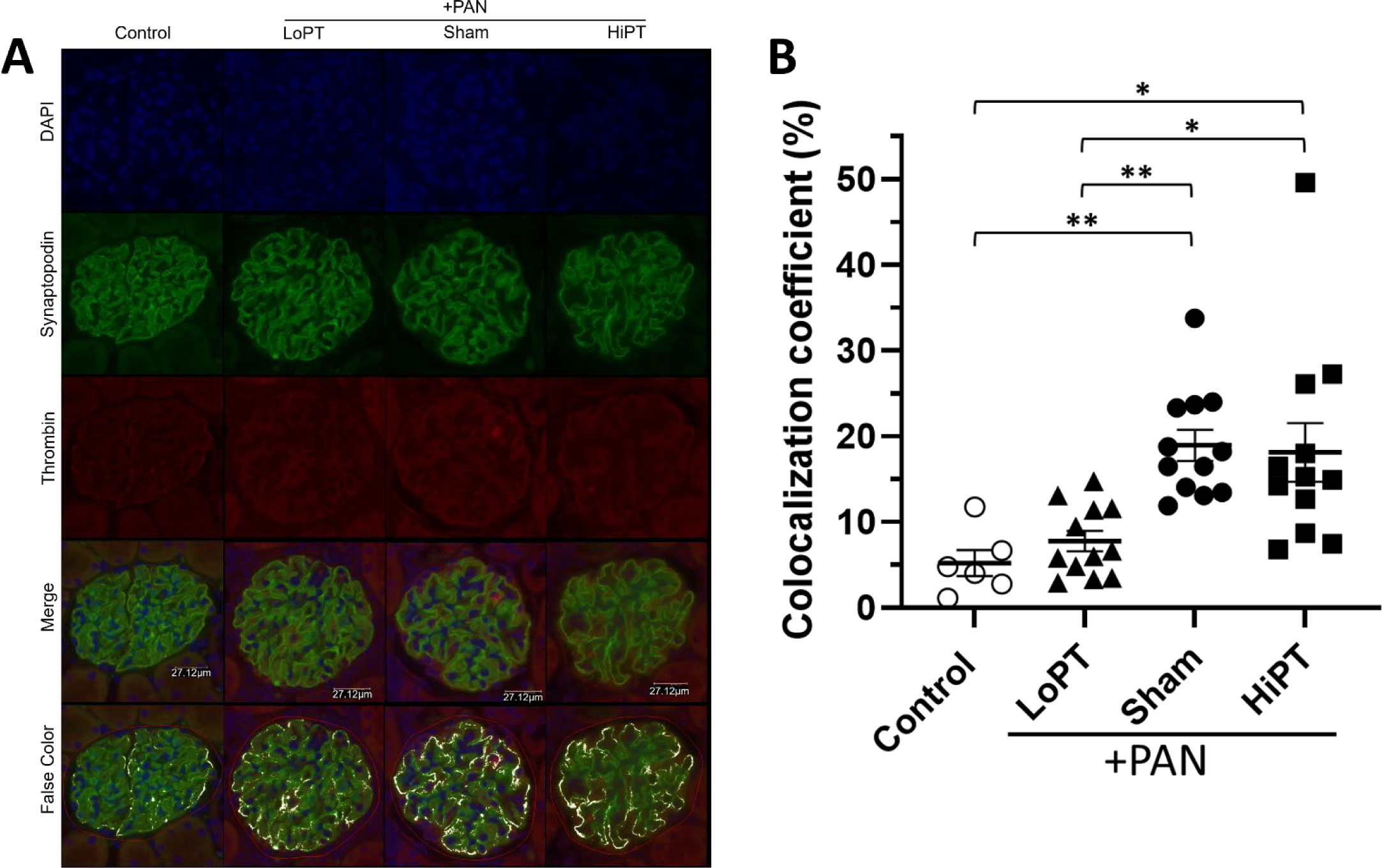
Plasma Prothrombin Levels Dictated Thrombin-Podocyte Interactions during Rat PAN-NS. (**A**) Representative immunofluorescence histology of glomeruli on day 10 (original magnification, 60x). (**B**) Relative proportion of synaptopodin-positive pixels (representing podocytes) with colocalized thrombin calculated using the colocalization coefficient feature of ZEN Black software (Zeiss USA). *n*=3-12 rats per group; each point represents averaged data taken from 20 random glomeruli per rat, \**P*<0.05, \*\**P*<0.01; ○: Control; ▴: LoPT (ASO-mediated hypo-prothrombinemia); ●: Sham; ▪: HiPT (prothrombin infusion-mediated hyper-prothrombinemia)

### Hypo-prothrombinemia Ameliorates Podocyte Foot Process Effacement

PAN-NS is known to cause podocyte foot process effacement and tubular epithelial injury (*10, 66, 67*). Directly relevant to the present study, high-dose antithrombin (a non-specific thrombin inhibitor) administration has been shown to reduce tubular injury in PAN-NS (*36, 68–71*). In the present study, renal cortex histology revealed tubular epithelial vacuolization consistent with mild acute tubular necrosis in PAN-NS (**Figure 5**). These findings were most pronounced in the HiPT group whereas the LoPT rats had normal histology. When a semiquantitative acute tubular necrosis score was applied, HiPT rats had significantly more tubular injury than LoPT rats (1.42±0.30 vs. 0.38±0.09; *P*<0.05). Importantly, no glomerular sclerosis was observed in any of the groups at this early PAN-NS timepoint. Electron microscopy revealed foot process effacement in all PAN-NS groups which was significantly reduced in the LoPT (44±7% vs. 76±4% in Sham; *P*<0.01; **Figure 5**). Additionally, fewer lipoprotein inclusions were observed in podocytes from LoPT rats as compared to Sham and HiPT. Importantly, we did not observe any mesangial, endothelial, or parietal epithelial abnormalities and there were no immune deposits in the glomerular basement membranes which had normal appearing ultrastructure. Collectively, these data strongly suggest that prothrombin knockdown reduces podocyte and tubular injury during PAN-NS.

**Figure 5:**
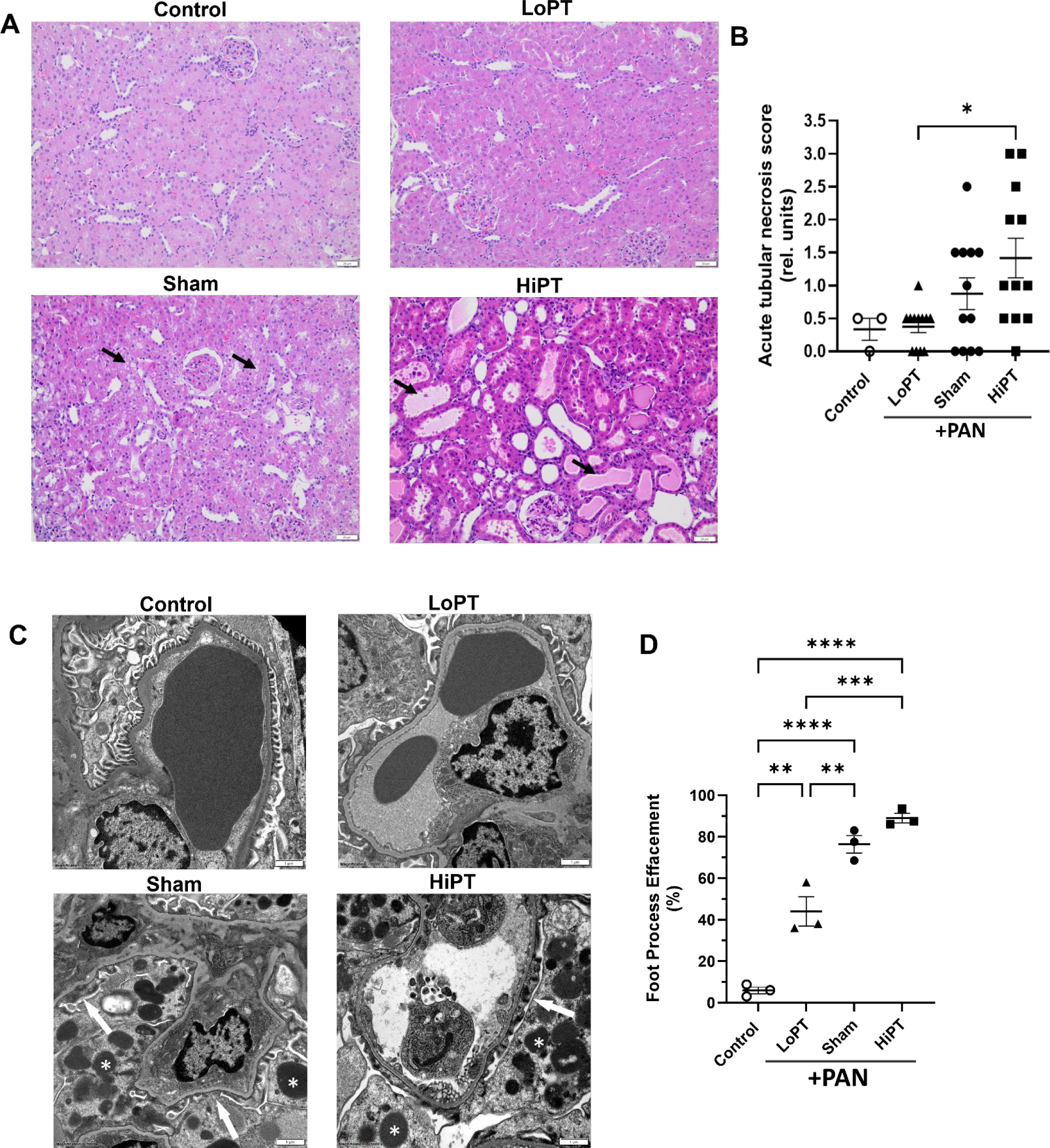
Hypo-prothrombinemia Ameliorates Podocyte Foot Process Effacement. (**A**) Representative photomicrographs of hematoxylin and eosin (H&E) stained renal cortex on day 10 (original magnification, 200x). Note vacuolization of proximal tubular epithelial cells consistent with acute tubular necrosis in the Sham image (arrows) and proximal tubular epithelial cell flattening and regenerative changes in the HiPT image (arrows). (**B**) Semiquantitative acute tubular necrosis score. *n*=3-12 rats per group. (**C**) Representative transmission electron microscopy images of renal cortex on day 10 (original magnification, 15,000x). Note podocyte foot process effacement (arrows) and lipoprotein inclusions (asterisks) in the Sham and HiPT images. (**D**) Percent foot process effacement. *n*=3 rats per group; each point represents averaged data taken from 20 glomerular capillary images per rat. \**P*<0.05, \*\**P*<0.01, \*\*\**P*<0.001; ○: Control; ▴: LoPT (ASO-mediated hypo-prothrombinemia); ●: Sham; ▪: HiPT (prothrombin infusion-mediated hyper-prothrombinemia)

### Hypo-prothrombinemia Diminished *in situ* Podocyte Injury

We and others previously demonstrated that *in vitro* thrombin-exposure leads to DNA nicking that is consistent with terminal podocyte injury (*34, 35*). Importantly, podocytes may be terminally injured prior to detachment from the glomerular capillary surface (*19*). Thus, to evaluate the health status of *in situ* podocytes we performed TUNEL assays on dissociated cells from isolated glomeruli. Importantly, neither LoPT nor HiPT altered podocyte DNA-nicking in otherwise healthy rats (**Figure S7**). However, HiPT significantly increased TUNEL-positive podocytes (17.4±0.5%) in PAN-NS (11.7±1.8% vs Sham; *P*<0.05; **Figure 6**). In contrast, LoPT reduced TUNEL-positivity (6.9±1.5%; *P*<0.05 vs. Sham) to levels not significantly different from Control (6.5±1.5%; *P*=0.90). Podocyte DNA-nicking was correlated with prothrombin levels, and thrombin-colocalization during PAN-NS (**Figure S8**).

**Figure 6:**
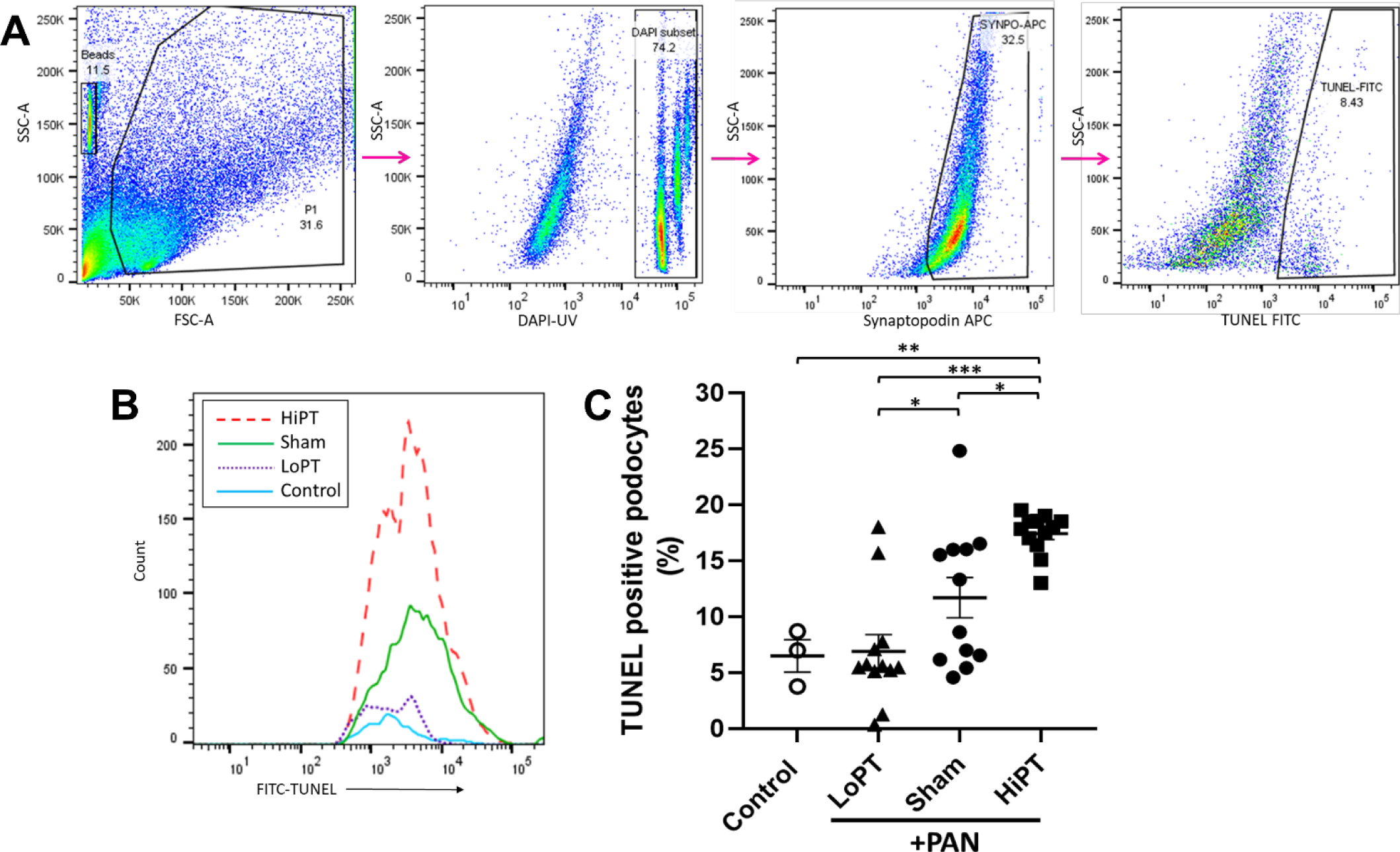
Hypo-prothrombinemia Diminished *in situ* Podocyte Injury. (**A**) Representative dot plot and subsequent gating strategy used to identify the proportion of synaptopodin-positive cells (podocytes) whose nuclei were TUNEL-positive, indicating DNA nicking (a marker of terminal podocyte injury). (**B**) Representative histograms of TUNEL-positive podocyte nuclei by treatment group. (**C**) On day 10, LoPT PAN-NS rats had significantly reduced TUNEL-positive podocyte nuclei, in contrast HiPT PAN-NS rats had significantly increased TUNEL-positive podocyte nuclei. *n*=3-12 rats per group; each point represents averaged data derived from ∼2,500 glomeruli and 100,000 event counts per rat, \**P*<0.05, \*\**P*<0.01, \*\*\**P*<0.001; SSC-A: side scatter-area; FSC-A: forward scatter-area; DAPI-UV: 4′,6-diamidino-2-phenylindole; Synaptopodin-APC: allophycocyanin-labeled anti-synaptopodin antibody; TUNEL-FITC: Fluorescein isothiocyanate-terminal deoxynucleotidyl transferase dUTP nick end labeling; ○: Control; ▴: LoPT (ASO-mediated hypo-prothrombinemia); ●: Sham; ▪: HiPT (prothrombin infusion-mediated hyper-prothrombinemia)

### Hyper-prothrombinemia Reduced *in situ* Podocyte Survival

Glomerular filtration barrier function and CKD progression are highly dependent on glomerular podocyte loss. We thus evaluated *in situ* podocyte survival by counting remaining podocytes in dissociated glomerular cells. Whereas LoPT and HiPT did not alter podocyte counts in otherwise healthy rats (**Figure S7**), HiPT PAN-NS rats had significantly lower podocyte counts (134.2±21.5 podocytes/glomerulus) vs. either LoPT (350.9±65.3; *P*<0.01) or Control (389.0±41.8; *P*<0.001; **Figure 7**). However, the LoPT and HiPT conditions did not translate into significantly improved or diminished podocyte counts vs. Sham. Meanwhile, podocyte counts were correlated with prothrombin levels, thrombin-colocalization, and podocyte injury in PAN-NS rats (**Figure S9**). These data suggest that hyper-prothrombinemia may accelerate podocyte loss during PAN-NS.

**Figure 7:**
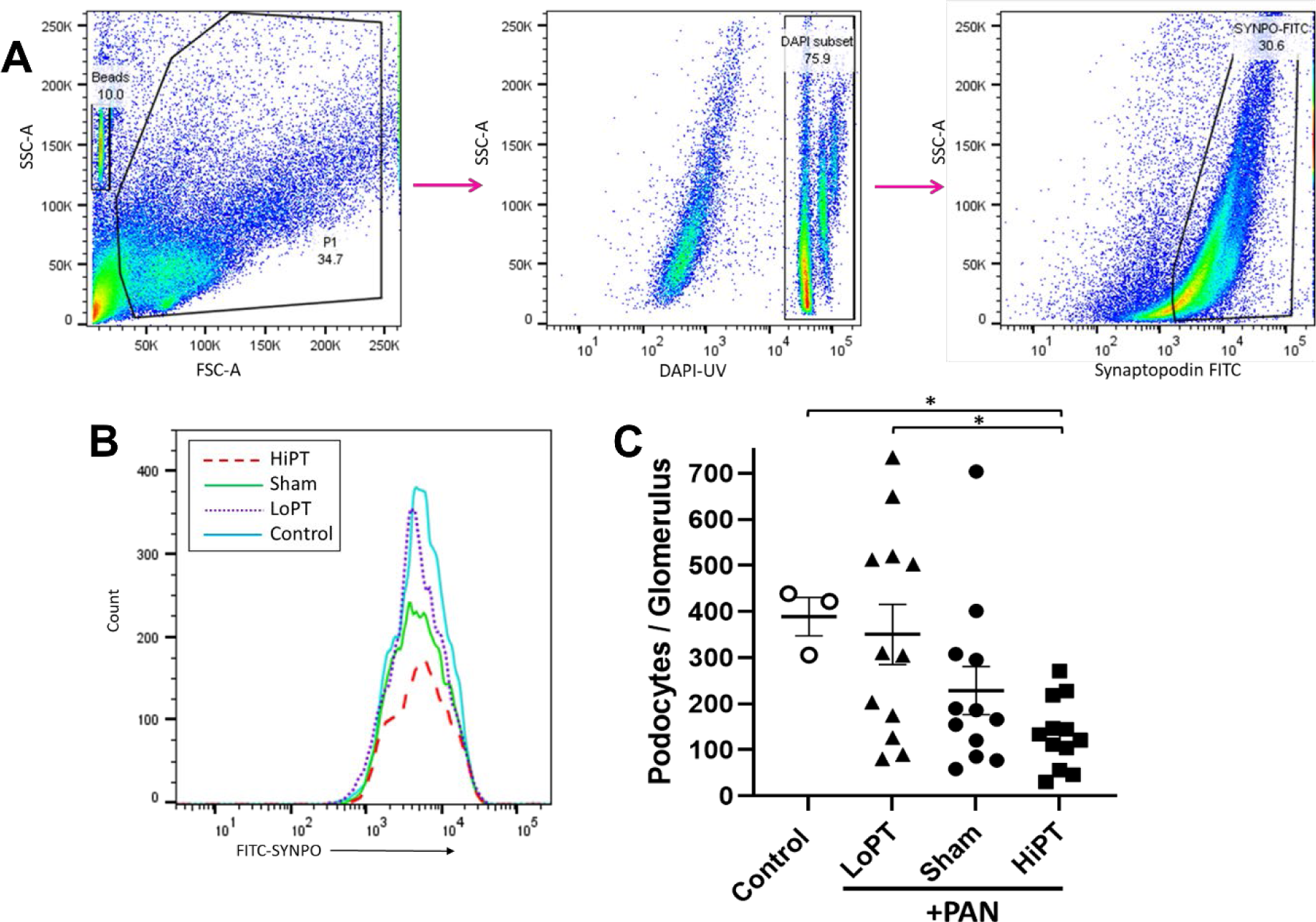
Hyper-prothrombinemia Reduced *in situ* Podocyte Survival. (**A**) Representative dot plot and subsequent gating strategy used to count synaptopodin-positive cells (podocytes). (**B**) Representative histograms of podocytes by treatment group. (**C**) On day 10, HiPT rats had significantly reduced podocyte counts in comparison to control and LoPT rats. *n*=3-12 rats per group; each point represents averaged data derived from ∼2,500 glomeruli and 100,000 event counts per rat; \**P*<0.05; SSC-A: side scatter-area; FSC-A: forward scatter-area; DAPI-UV: 4′,6-diamidino-2-phenylindole; Synaptopodin-FITC: Fluorescein isothiocyanate-labeled synaptopodin; ○: Control; ▴: LoPT (ASO-mediated hypo-prothrombinemia); ●: Sham;: HiPT (prothrombin infusion-mediated hyper-prothrombinemia)

### Prothrombin Regulated Proteinuria and Plasma Albumin during Puromycin Aminonucleoside-Induced Rat Nephrotic Syndrome

Podocyte injury and loss leads to glomerular filtration barrier dysfunction resulting in proteinuria and hypoalbuminemia (*10, 16, 18, 20*). Consistent with preserved podocyte function, LoPT PAN-NS rats had partially improved day 10 proteinuria values that were significantly different from Sham (6.6±0.7 vs. 15.0±3.7 mg/mg creatinine; *P*<0.05) but were not significantly different than Control (1.0±0.1 mg/mg; *P*=0.56; **Figure 8**). Meanwhile, HiPT rats had significantly worse proteinuria (34.9±9.3 mg/mg; *P*<0.05). Consistent with improved overall protein homeostasis, plasma albumin was significantly improved in LoPT vs. Sham (3.32±0.03 vs. 3.07±0.08 g/dL; *P*<0.05) to values that were not significantly different from Control (3.41±0.02 g/dL). However, HiPT did not significantly worsen hypoalbuminemia (3.05±0.1 g/dL). Importantly, neither LoPT nor HiPT altered proteinuria or plasma albumin in otherwise healthy rats (**Figure S7**). In PAN-NS proteinuria was significantly correlated with prothrombin levels and podocyte injury (**Figure 8**). Similarly, plasma albumin was correlated with prothrombin levels, podocyte injury, podocyte survival, and proteinuria (**Figure S10**).

**Figure 8:**
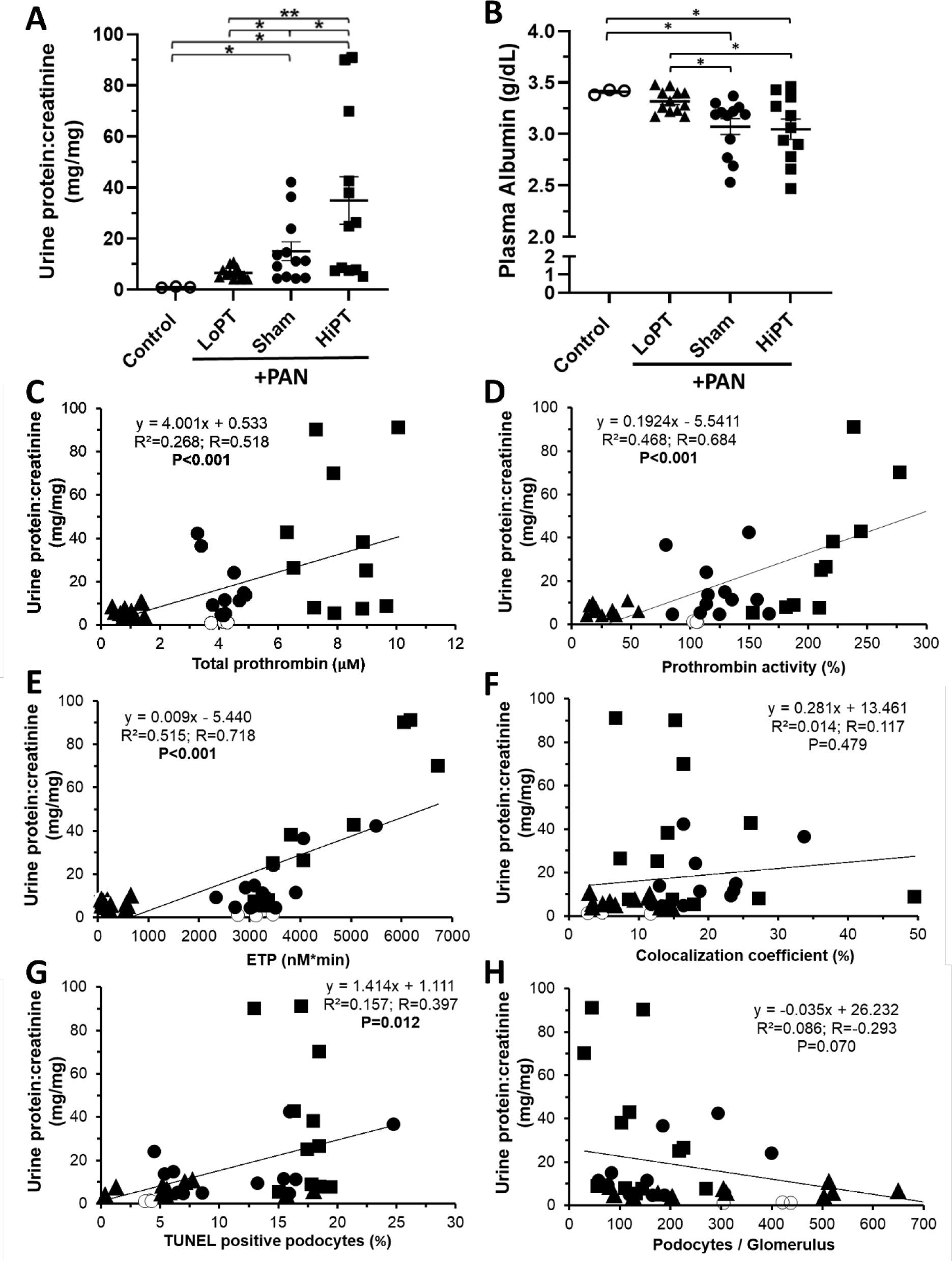
Prothrombin Regulated Proteinuria and Plasma Albumin during Rat PAN-NS. (**A**) Day 10 proteinuria was significantly reduced in LoPT rats whereas HiPT rats had significantly more proteinuria than any other group. (**B**) Plasma albumin was significantly improved in LoPT rats. Proteinuria was significantly correlated with plasma prothrombin protein quantity (**C**), enzymatic activity (**D**), and endogenous thrombin potential (ETP, **E**). Proteinuria correlation with podocyte-prothrombin colocalization, TUNEL-positive podocytes, and podocyte counts per glomerulus are shown in panels **F-H**, respectively. *n*=3-12 rats per group; \**P*<0.05, \*\**P*<0.01; ○: Control; ▴: LoPT (ASO-mediated hypo-prothrombinemia); ●: Sham; ▪: HiPT (prothrombin infusion-mediated hyper-prothrombinemia)

## DISCUSSION

Proteinuria, podocytopathy, and podocytopenia all contribute to glomerular dysfunction and progression of chronic kidney disease (*8, 16, 22, 72–78*). The data presented in this study demonstrate for the first time that circulating levels of prothrombin (the zymogen precursor of thrombin) modulate *in vivo* podocyte thrombin exposure, injury, survival, and function in the setting of proteinuric glomerular disease. We and others have previously shown that thrombin injures cultured podocytes *in vitro* and that anticoagulant-mediated thrombin inhibition preserves *in vivo* glomerular function as determined by proteinuria reduction (*34–38*). In previous studies on this topic, the anticoagulant agents may have reduced proteinuria via alternate mechanisms (i.e. upstream coagulation factor inhibition, effects on non-podocyte cells) (*79, 80*). In contrast, this study directly manipulated prothrombin levels *in vivo*, providing direct evidence that (pro)thrombin is a driver of podocytopathy during proteinuria. We previously demonstrated that thrombin injures *in vitro* podocytes via its cognate protease-activated receptors (*34*). Collectively, these data suggest that thrombin originates as circulating prothrombin that is pathologically filtered during proteinuria leading to intraglomerular thrombin generation and direct podocytopathic thrombin signaling. Thus, interruption of this signaling pathway may represent a novel therapeutic target to slow or halt glomerular disease progression toward CKD.

Mounting indirect evidence from *in vitro* studies and *in vivo* pharmacologic manipulation strongly suggest that thrombin is a pathological driver of podocyte injury in the setting of proteinuria (*34–38*). The present study provides direct evidence supporting this hypothesis. HiPT rats exhibiting prothrombin levels ∼195% of normal developed worse proteinuria along with increased podocyte injury. The human thrombophilic single nucleotide polymorphism, *F2* G20210A (rs1799963), is associated with prothrombin levels ∼115-170% of normal (*49*). To our knowledge, rs1799963 has not been associated with enhanced CKD progression or incidence, but most studies were not directly looking for such a signal, were likely underpowered, or were investigating CKD-related thrombotic risk (*81–83*). It is also possible that since the reference interval for prothrombin extends to 150% that the mildly increased levels associated with this polymorphism do not lead to meaningfully increased CKD risk (*84*). However, the present data suggest that rs1799963 should be more carefully analyzed as a potential driver of proteinuria-mediated CKD progression. Protein overload, achieved with ≥3 g/kg cumulative intravenous albumin, is a commonly employed method to induce transient proteinuria in animal models (*10, 85*). However, it is unlikely that HiPT mimics protein overload since the cumulative prothrombin protein dose (81.26 mg/kg) used in these studies is only 2.7% of the minimum albumin dose required to induce proteinuria and did not induce proteinuria in the absence of PAN (**Figure S7**).

In contrast, highly specific ASO-mediated prothrombin knockdown (to ∼24% of normal) resulted in diminished podocyte thrombin exposure, podocytopathy, podocytopenia, and proteinuria with improved plasma albumin. Prothrombin is predominantly synthesized in the liver but several other organs, including the kidney, also produce prothrombin (*62*). Moreover, ASO therapy has been demonstrated to reduce target mRNA in many tissues, including both liver and kidney (*86*). Thus, while it is clear from these experiments that ASO-mediated prothrombin knockdown improves podocyte function and survival, the origin of podocytopathic prothrombin remains ill-defined. However, manipulation of circulating prothrombin did not induce proteinuria in healthy animals, strongly suggesting that proteinuria is a prerequisite for (pro)thrombin-mediated podocyte injury and that therefore podocytopathic prothrombin most likely originates from the plasma compartment. Alternatively, podocyte injury or proteinuria may stimulate glomerular cells or adjacent tissues to upregulate local prothrombin synthesis. A previous study identified prothrombin protein in tubular epithelial cells, but not glomeruli, suggesting that juxtaglomerular tubules could be a relevant local source of prothrombin synthesis (*62*).

We previously demonstrated that *in vitro* thrombin-mediated podocytopathy is dependent upon activation of podocyte-expressed protease-activated receptors (*34*). Activation of these receptors is dependent upon enzymatic cleavage by thrombin or other enzymes resulting in exposure of a cryptic tethered ligand (*87*). Once generated, thrombin has a <1 minute half-life *in vivo* (*84, 88–90*), suggesting that prothrombin is most likely converted to thrombin locally in the glomerulus, perhaps even on the podocyte surface (*88, 91*). However, the mechanism by which prothrombin is converted to intraglomerular thrombin during proteinuria remains undefined. The canonical hemostatic mechanism is prothrombin cleavage by the factor Xa/Va prothrombinase complex. While prothrombin and cofactor V have been found in the urine of nephrotic patients, factor X has not (*92, 93*). Additionally, a proteomic study has demonstrated that neither factor X nor V protein are expressed by healthy murine podocytes (*94*). Thus, while podocytes do express tissue factor, the complete complement of canonical prothrombinase components do not appear to be available (*91, 95, 96*). Alternatively, adjacent kidney cells have been reported to synthesize both factor X and V (*97*). Moreover, these components may originate from the plasma during proteinuria. It is thus possible that the canonical prothrombinase mechanism is at work in the diseased glomerulus. Alternatively, one or more putative tissue prothrombinases may explain glomerular thrombin activity (*98*). Amongst the latter, fibrinogen-like protein 2 (*Fgl2*) is expressed by podocytes and is thus a leading candidate (*94, 99–101*). This possibility is particularly intriguing as it would provide a potential target to ameliorate (pro)thrombin-mediated podocytopathy without increasing the risk of CKD-related bleeding (*102*).

No spontaneous bleeding or thrombotic events were observed in these experiments, suggesting that thrombin inhibition may be well-tolerated during nephrotic syndrome. Indeed, anticoagulant prophylaxis is a recommended intervention in the management of glomerular disease and these data suggest that this practice should be studied for potential antiproteinuric benefits (*40*). Although hyper-prothrombinemia reduced podocyte counts to 38% of healthy control values, no glomerulosclerosis was observed in these acute PAN-NS studies. Additional studies using chronic NS models will be needed to fully elucidate the effects of (pro)thrombin levels on the evolution of focal segmental glomerulosclerosis and CKD progression. Interestingly, thrombin-podocyte colocalization did not increase in the setting of hyper-prothrombinemia. This somewhat unexpected observation may indicate that podocyte-expressed thrombin receptors are already saturated at normal prothrombin concentrations during proteinuria (*34*). Alternatively, thrombin-mediated podocyte injury earlier in the experimental timeline may have downregulated protease-activated receptor expression. The area under the thrombin-time exposure curve is known to influence endothelial responses (*103*). Thus, the podocyte response may be dependent on both concentration and time which may explain worse outcomes during hyper-prothrombinemia without an increase in thrombin-podocyte colocalization. Tubular epithelial injury is a known feature of PAN-NS that is proportional to proteinuria (*10, 66, 67*). We observed reduced proximal tubular injury in the setting of hypo-prothrombinemia, which is consistent with the previous observations that reduced proteinuria coincides with ameliorated tubular injury. Alternatively, it is possible that (pro)thrombin may directly exert cytotoxic effects on tubular epithelium or activate resident interstitial inflammatory cells and pathways during proteinuria (*80*).

There are several limitations to consider when interpreting these experiments. Although the mammalian coagulation system is highly conserved, it is possible that the use of human prothrombin in these studies could have resulted in subtle species-dependent differences in prothrombin activation efficiency or protease-activated receptor signaling responses in the rat glomerulus. The persistent hyper-prothrombinemia and lack of glomerular immune deposits after multiple human prothrombin injections argues against both xenoimmunity in this model and an immune-mediated mechanism for worse outcomes in the HiPT rats. Nonetheless, the use of rat prothrombin or creation of a gene duplication-mediated hyper-prothrombinemia model would provide more specific evidence. Although PAN is a relatively podocyte-specific toxin, it is possible that there are off-target PAN effects on the coagulation system. However, PAN reportedly does not alter protein synthesis in isolated rat hepatocytes, making it unlikely that PAN directly altered coagulation protein synthesis (*104*). We demonstrated significant correlation between histologic- and flow cytometry-based podocyte counting methods. However, the limitations of histological podocyte counting are well-described and the method used in these studies did not have adequate sensitivity to detect between group differences in these acute PAN-NS experiments (*53*). This may partly be explained by the infeasibility of manually counting hundreds or thousands of glomeruli per animal in a high-throughput manner and because glomeruli that are larger in diameter than the microscope objective’s field of view cannot be easily captured, both issues were overcome by the flow cytometry approach. While it is possible that the digestion buffer used to dissociate the glomeruli for flow cytometry analysis may have injured the podocytes and thereby increased the *ex vivo* TUNEL signal. Therefore, this method may have systematically overestimated *in situ* TUNEL positivity. Nonetheless, all samples were handled identically and thus the relative differences between the groups are likely relevant, especially since they correlate well with all of the other NS parameters. Finally, these data, generated in a rat NS model, may not be generalizable to other forms of glomerular disease that lead to CKD such as diabetic nephropathy, lupus nephritis, or hypertensive nephropathy and thus, should be examined in animal models of these glomerular diseases.

In summary, prothrombin modulates both podocyte function and survival in the PAN model of nephrotic syndrome. To determine the importance of prothrombin-mediated podocyte injury and its potential as a novel therapeutic target to slow CKD progression will require additional studies using a chronic NS model. Important next steps include defining the relevance of this mechanism in CKD models, determining the prothrombinase mechanism driving intraglomerular thrombin formation, and evaluating its suitability as a druggable target using thrombin inhibitors, especially those that are already approved by relevant regulatory agencies for use in humans with kidney disease and may thus be suitable for rapid clinical translation (*79*). Because proteinuria imparts increased thrombotic risk (*29, 31, 105–107*), renally excreted anticoagulant agents that may inhibit intraglomerular thrombin should be investigated as a novel means to simultaneously reduce thrombotic complication risk and slow proteinuria-mediated CKD progression.

## Acknowledgments

This project was supported by grants K08DK103982, R03DK118315, and R01DK124549 (to BAK) from the National Institute of Diabetes and Digestive and Kidney Diseases (NIDDK). The content is solely the responsibility of the authors and does not necessarily represent the official views of the National Institutes of Health. Additional support was provided by the George & Elizabeth Kelly Foundation (Lewis Center, OH; to BAK). This study was also supported (in part) by research funding from the CSL Behring – Professor Heimburger Award to BAK. The authors are indebted to Dr. Brett P. Monia, Ionis Pharmaceuticals, for providing the antisense oligonucleotides, Matthew Curtis, Carl Zeiss Microscopy, for developing the Zen Blue software algorithm for histological podocyte counting, and Dave Dunaway, Abigail Wexner Research Center, for assistance with flow cytometry protocol refinement.

## Authorship Contributions

Experiments were conducted by APW, KJW, IP, ZSS, EA, KM, TKW, ARB, EPC, TAV, SVB, and BAK. APW and BAK analyzed data; APW and BAK prepared the figures and wrote the paper; all authors read, edited, and approved the final version; WES and MTN provided important intellectual contributions; BAK was responsible for funding, oversight, and coordination of the project.

## Conflict of Interest Disclosures

The authors declare no competing financial interests.

## Notes

**Support:** This project was supported by grants K08DK103982, R03DK118315, and R01DK124549 (to BAK) from the National Institute of Diabetes and Digestive and Kidney Diseases (NIDDK). The content is solely the responsibility of the authors and does not necessarily represent the official views of the National Institutes of Health. Additional support was provided by the George & Elizabeth Kelly Foundation (Lewis Center, OH; to BAK). This study was also supported (in part) by research funding from the CSL Behring – Professor Heimburger Award to BAK.

### Competing Interest Statement

The authors have declared no competing interest.

### Summary of Updates

Additional data added: new Figure 5, revised Figure S3, new Movies M1-M4. Minor revisions to text. Authors updated.

