## SUPPLEMENTAL METHODS AND DATA for "Prothrombin Knockdown Protects Podocytes and Reduces Proteinuria in Glomerular Disease"

#### Prothrombin Drives Podocyte Injury and Enhances Proteinuria in Glomerular Disease

### SUPPLEMENTAL METHODS

#### Reagents and Resources

Antisense oligonucleotides 401027 (TGTGATTTTGCAATGAGGAT; 100% complementary to both mouse and rat *F2* mRNA), 401025 (ATTCCATAGTGTAGGTCCTT; 100% complementary to mouse *F2* mRNA, 75% to rat *F2* mRNA), and 141923 (CCTTCCCTGAAGGTTCCTCC; scrambled control) were a kind gift from Ionis Pharmaceuticals, Carlsbad, CA. Human prothrombin was from Prolytix (Essex Junction, VT). Digestion Buffer: Dispase II (2.8 units/mL; Sigma-Aldrich, St. Louis, MO), type 4 collagenase (300 units/mL) and DNase I (50 units/mL; both from Worthington Biochemical Corp., Lakewood, NJ) in HBSS (ThermoFisher, Waltham, MA). Antibodies were fluorescently conjugated using FITC and APC kits from Abcam (Cambridge, MA), as indicated. Mouse monoclonal synaptopodin antibody (10R-S125A) was purchased from Fitzgerald Industries International (Acton, MA, USA). Rabbit polyclonal thrombin antibody (bs-1914R; raised against the thrombin heavy chain) was from Bioss (Woburn, MA). Secondary antibodies (Alexa Fluor 488 anti-mouse IgG and Alexa Fluor 594 antirabbit IgG) were from ThermoFisher. See Table S2 for additional reagents and resources.

#### Cell Culture

Primary rat hepatocytes from male Wistar rats were from Life Technologies (Waltham, MA). In brief, rat hepatocytes were grown on plates coated with collagen type 1 at 37°C in Williams' E medium (Invitrogen, Waltham, MA) with 5% Fetal Bovine Serum (Fisher, USA). For experiments, hepatocytes were seeded at $10^6$ cells/well in 6-well plates and allowed to expand to ~80% confluence over 48 hours. Cells were washed with PBS and then incubated with media plus antisense oligonucleotide (control ASO, ASO 401025, or ASO 401027) at 5 µM. After 72 hours, the cells were washed with PBS and lysed with RLT buffer (Qiagen, Valencia, CA) containing 1% β-Mercaptoethanol (*1*).

#### Healthy Control Animals

Hypo- (LoPT) and hyper-prothrombinemia (HiPT) was modeled without PAN to exclude podocyte effects in the absence of proteinuria. LoPT rats received anti-*F2* ASO (401027) on Days -17.5 to +9 and saline (instead of PAN) on Day 0. HiPT rats received prothrombin infusions on Days 0 to +9 and saline (instead of PAN) on Day 0.

**RNA Extraction and Reverse Transcriptase – Quantitative Polymerase Chain Reaction**

Whole rat liver was stored at -80C in RNAlater (ThermoFisher, Waltham, MA) until lysed in RLT buffer (Qiagen,Valencia, CA) containing 1% β-Mercaptoethanol (*1*). Total RNA was isolated from primary rat hepatocytes or whole rat liver using the RNeasy kit (Qiagen, USA). cDNA was made from 0.5 µg total RNA using the Protoscript First Strand cDNA Synthesis kit (New England Biolabs; Ipswich, MA), according to the manufacturer's instructions. Purity and yield of RNA and cDNA were confirmed by measuring the absorbance at 260 and 280 nm. cDNA was amplified by quantitative polymerase chain reaction (qPCR); 0.5 µL cDNA (diluted in a total volume of 11 µL RNase-free H2O), 0.5 uL forward (fwd) and reverse (rvs) primers (below), and 12.5 uL iQ SYBR Green Supermix (Bio-Rad, Hercules, CA). Primers were obtained from Invitrogen (Carlsbad, CA): β-actin (*Actb*, fwd: CGGTCCACACCCGCCACC, rvs: CTTGCTCTGGGCCTCGTCGC) and Prothrombin (*F2*, fwd: GGACGCTGAGAAGGGTATCG, rvs: GGTGGTTGTAGGGGCTCTTC). qPCR was performed as follows: 1 cycle at 95 °C for 5 min, 40 cycles (melting at 94 °C for 30 sec; annealing gradient 55→72 °C for 30 sec; extension at 72 °C for 30 sec), with a final extension of 72 °C for 10 min using a CFX96 Real-Time System thermal cycler (Bio-Rad, Hercules, CA). All RT-qPCR samples were run in triplicate and reported as mean *F2* expression relative to mean housekeeping gene (*Actb*) expression.

**Immunofluorescent Three-Dimensional Podocyte Counts**

Aliquots of the glomerular suspensions (3000-6000 glomeruli) were prepared for whole mount immunofluorescence histology using an adaptation of previously reported methods for whole mount analysis of *D. melanogaster* organs (*2, 3*). Glomeruli were fixed in 4% paraformaldehyde for 35 minutes, washed in Incubation Buffer (0.5% Triton X-100 in PBS supplemented with 5% FBS) for 10 minutes x3, then stored in 70% ethanol at -20ºC until ready for analysis. The glomeruli were then washed once in Incubation Buffer, transferred to an optically pure 96 well plate, sealed with an evaporation barrier, and photobleached as described in

Methods, Immunofluorescence Histology. The glomeruli were then transferred to 600 µL Eppendorf tubes, permeabilized with 1% Triton X-100 in PBS supplemented with 5% FBS for 60 minutes, washed x3 in Incubation Buffer, and blocked with 0.5% Triton X-100 in SuperBlock for 30 minutes. Glomeruli were then stained with anti-WT-1 primary antibody (mouse IgG1 (Novus Biologicals, Centennial, CO) at 1:200 in Incubation Buffer) at 4ºC for 4 days on a vertical rotator, washed in Incubation Buffer x3, followed by secondary anti-mouse IgG1 (Invitrogen, Waltham, MA) at 4ºC for 4 days, washed in Incubation Buffer x3, then equilibrated in 30 µL Prolong Gold with DAPI (Invitrogen) for 30 minutes. Stained glomeruli (15 µL; 1500-3000 glomeruli) were then whole mounted on glass slides under a coverslip. Random glomeruli (*n*=15/rat) were imaged as 1 µm z-stacks with a resolution of 0.391 µm/pixel using a 63x Plan Apochromat objective on a Zeiss LSM 700 confocal microscope (Carl Zeiss Microscopy, LLC, White Plains, NY) with the 405 nm and 488 nm lasers and detection of DAPI and AlexaFluor 488, respectively. Z-stacks were analyzed in 3D volume for WT-1-positive nuclei using NIS-Elements software version 5.30 (Nikon Instruments, Melville, NY). Z-stacks were preprocessed with Denoise.ai to facilitate segmentation, and 3D bright spot detection was used on the DAPI channel to automatically identify nuclei (*4, 5*). The centroid of each identified DAPI nucleus was marked with a 3D spot with a 3-voxel diameter. An automated threshold was calculated from the WT-1 channel intensity in each Z-stack and used to segment WT-1-positive voxels, and any DAPI centroid intersecting WT-1-positive signal was marked as a WT-1-positive nucleus, similarly to a previously published analysis method (*6*).

**Histology and Electron Microscopy**

Formalin fixed paraffin embedded renal cortex from each rat was sectioned (3 µm) and mounted on glass slides using standard techniques. After hematoxylin and eosin staining, each section was examined and scored by a renal pathologist (SVB) who was blinded to the study group. Electron microscopy analysis was performed on the 3 rats nearest to the mean proteinuria from each group. Optimal cutting temperature (OCT) embedded renal cortex was cut into 1 $mm^3$ pieces and fixed in 3% glutaraldehyde and stored at 4ºC until further use. Subsequently, samples were washed in 0.2M Sodium Cacodylate buffer x2 for 10 minutes then post-fixed in 1% osmium tetroxide for 1 hour. Samples were rinsed in sym-collidine buffer x2 for 10 minutes followed by *en bloc* staining in 2.5% uranyl acetate solution. Samples were dehydrated using a graded ethanol series (30 – 95%), followed by 3 rinses of 100% ethanol and 3 rinses of acetone. Samples were placed in diluted Spurr's resin solution (2 parts resin to 1 part acetone) overnight. The following morning specimens were embedded in 100% Spurr's resin (EMS). Blocks were polymerized overnight at 65°C before cutting on a Leica U6 ultramicrotome. Thick sections were cut at 750 nm and stained with methylene blue/Azur II at 80°C for 10 minutes and counter-stained with basic fuchsin at 60°C for 23 seconds. Thick sections were evaluated, and blocks were trimmed down to regions of interest. Thin sections (90 nm) were collected on 200 mesh Cu grids for electron microscopy evaluation. Specimens were evaluated on a JEOL JEM-1400 transmission electron microscope (TEM) operating at 80 kV and images were collected on an Olympus Veleta digital camera using Radius software. The TEM images were then examined and scored by an investigator (IP) under the supervision of a renal pathologist (SVB), both of whom were blinded to the study group.

**Other**

Plasma albumin concentrations were determined using bromocresol purple (BCP) assay (QuantiChrom BCP; BioAssay Systems, Hayward, CA), as described previously (*7, 8*). Proteinuria was determined by urinary protein-to-creatinine (UPC) ratio both of which were quantified by Antech Diagnostics (Morrisville, NC), using standard techniques that are fully compliant with Good Laboratory Practice regulations, as previously reported (*1, 7, 9, 10*).

**Table S1: ARRIVE Guidelines for Reporting *In Vivo* Animal Experiments**

| | **Dose-finding Preliminary Study Rats** | **Main Study Nephrotic Rats#** | **Main Study Healthy Control Rats#** |
|---|---|---|---|
| **Total Rats** | N=24 Wistar rats males | N=36 Wistar rats males | N=9 Wistar rats males |
| **PAN dose** | None | 50 mg/kg, in 500 uL sterile saline; IV; once | None |
| **Treatment Groups (N/group)** | i) LoPT= PT-ASO @ 0, 40, 50, or 60 mg/kg, in 500 uL sterile saline; IP; twice weekly for 17.5 days (2/dose→N=8)<br>ii) LoPT= PT-ASO @ 60 mg/kg, in 500 uL sterile saline; IP; twice weekly for 27.5 days (N=3)<br>iii) HiPT= hPT @ 31.25 mg/kg, in 500 uL sterile saline; IV; once (2/timepoint→N=10)<br>iv) HiPT= hPT @ 31.25 mg/kg, in 500 uL sterile saline; IV; on Day 0 +16.67 mg/kg on Days 3, 6, & 9 (N=3) | i) LoPT= PT-ASO @ 60 mg/kg, in 500 uL sterile saline; IP; twice weekly for 27.5 days (N=12)<br>ii) Sham= scrambled-ASO @ 60mg/kg, in 500 uL sterile saline; IP; twice weekly for 27.5 days (N=12)<br>iii) HiPT= hPT @ 31.25 mg/kg, in 500 uL sterile saline; IV; on Day 0 +16.67 mg/kg on Days 3, 6, & 9 (N=12) | i) LoPT= PT-ASO @ 60 mg/kg, in 500 uL sterile saline; IP; twice weekly for 27.5 days (N=3)<br>ii) Control= scrambled-ASO @ 60mg/kg, in 500 uL sterile saline; IP; twice weekly for 27.5 days (N=3)<br>iii) HiPT= hPT @ 31.25 mg/kg, in 500 uL sterile saline; IV; on Day 0 +16.67 mg/kg on Days 3, 6, & 9 (N=3) |
| **Manuscript** | Figures 1 & 2 | Figures 3, 4, 5, 6, 7, S2, S3, S6, S8, S9 & S10 | Figures S6 & S7 |
| **ARRIVE Guidelines** | Randomized groups; Investigators performed sample collection and assays blinded | | |

#Note: all rats were subjected to identical experimental procedures, such that each group received their respective treatments plus corresponding sham saline injections to match the other two groups. Specifically: LoPT rats received 500 uL IV sham saline injections on Days 0, 3, 6, & 9. HiPT rats received 500 uL IP sham saline injections twice weekly for 27.5 days. Sham rats received 500 uL IV sham saline injections on Days 0, 3, 6, & 9, and IP injections of a scrambled control antisense oligonucleotide in 500 uL sterile saline twice weekly for 27.5 days.

IP: intraperitoneal, IV: intravenous, PT: prothrombin, hPT: human prothrombin, ASO: antisense oligonucleotide

### Table S2: Reagents & Resources

| Reagent Name | Company | Company Location |
|---|---|---|
| Antisense oligonucleotide 401027 | Ionis Pharmaceuticals | Carlsbad, CA |
| Antisense oligonucleotide 401025 | Ionis Pharmaceuticals | Carlsbad, CA |
| Antisense oligonucleotide 141923 | Ionis Pharmaceuticals | Carlsbad, CA |
| Human Prothrombin | Prolytix | Essex Junction, VT |
| Digestion buffer: dispase II | Sigma-Aldrich | St. Louis, MO |
| Type 4 collagenase | Worthington Biochemical Corp | Lakewood, NJ |
| DNase I | Worthington Biochemical Corp | Lakewood, NJ |
| HBSS | Thermofisher | Waltham, MA |
| FITC Conjugation Kit (1mg) | Abcam | Cambridge, MA |
| APC Conjugation Kit (1mg) | Abcam | Cambridge, MA |
| Mouse monoclonal synaptopodin antibody (10R-S125A) | Fitzgerald Industries International | Acton, MA |
| Rabbit polyclonal thrombin antibody (bs-1914R) | Bioss | Woburn, MA |
| Secondary antibody (Alexa Fluor 488 anti-mouse IgG) | Thermofisher | Waltham, MA |
| Secondary antibody (Alexa Fluor 594 anti-rabbit IgG) | Thermofisher | Waltham, MA |
| Primary rat hepatocytes | Life technologies | Waltham, MA |
| Williams' E medium | Invitrogen | Waltham, MA |
| 5% fetal bovine serum | Fisher | USA |
| RLT buffer | Qiagen | Valencia, CA |
| RNAlater | Thermofisher | Waltham, MA |
| RNeasy kit | Qiagen | USA |
| Protoscript First Strand cDNA synthesis kit | New England Biolabs | Ipswich, MA |
| iQ SYBR Green Supermix | BioRad | Hercules, CA |
| Primer β-actin (Actb, fwd: CGGTCCACACCCGCCACC) | Invitrogen | Carlsbad, CA |
| Primer β-actin (rvs: CTTGCTCTGGGCCTCGTCGC) | Invitrogen | Carlsbad, CA |
| Primer prothrombin (F2) | Invitrogen | Carlsbad, CA |
| Primer prothrombin (fwd: GGACGCTGAGAAGGGTATCG) | Invitrogen | Carlsbad, CA |
| Primer prothrombin (rvs: GGTGGTTGTAGGGGCTCTTC) | Invitrogen | Carlsbad, CA |
| CFX96 Real-time system thermal cycler | biorad | Hercules, CA |
| Anti-WT-1 primary antibody- mouse IgG1 | Novus Biologicals | centennial, CO |
| Secondary anti-mouse IgG1 | Invitrogen | Carlsbad, CA |
| Prolong gold with DAPI | Invitrogen | Carlsbad, CA |
| Zeiss LSM 700 microscope | Carl Zeiss Microscopy, LLC | White Plains, NY |
| ZEN blue | Zeiss USA | Thornwood, NY |
| Quantichrom BCP assay kit | Bioassay systems | Hayward, CA |
| Urine protein:creatinine measurement | Antech diagnostics | Morrisville, NC |
| Puromycin aminonucleoside | MilliporeSigma | Burliington, MA |
| Corn Trypsin Inhibitor | CTI, Prolytix | Essex Junction, VT |
| Live/dead near IR dye and FITC-conjugated synaptopodin antibody | Fitzgerald | North Acton, MA |
| LSR II flow cytometer | BD biosciences | Franklin Lakes, NJ |
| Leica DMI 4000B inverted fluorescence microscope | Leica | Deerfield, IL |
| ZEN Black software | Zeiss USA | Thornwood, NY |
| Rat and human-sepcific ELISA | MyBioSource Inc | San Diego, CA |

| | | |
|---|---|---|
| Rox prothrombin | DiaPharma | West Chester, OH |
| Technothrombin TGA kit | Technoclone | Vienna, Austria |
| Spectramax M2 fluorescent plate reader | Molecular devices | Sunnyvale, CA |
| SigmaStat software | Systat | San Jose, CA |
| GraphPad Prism | GraphPad Software Inc | La Jolla, CA |
| NIS-Elements | Nikon Instruments | Melville, NY |

**SUPPLEMENTAL FIGURES**

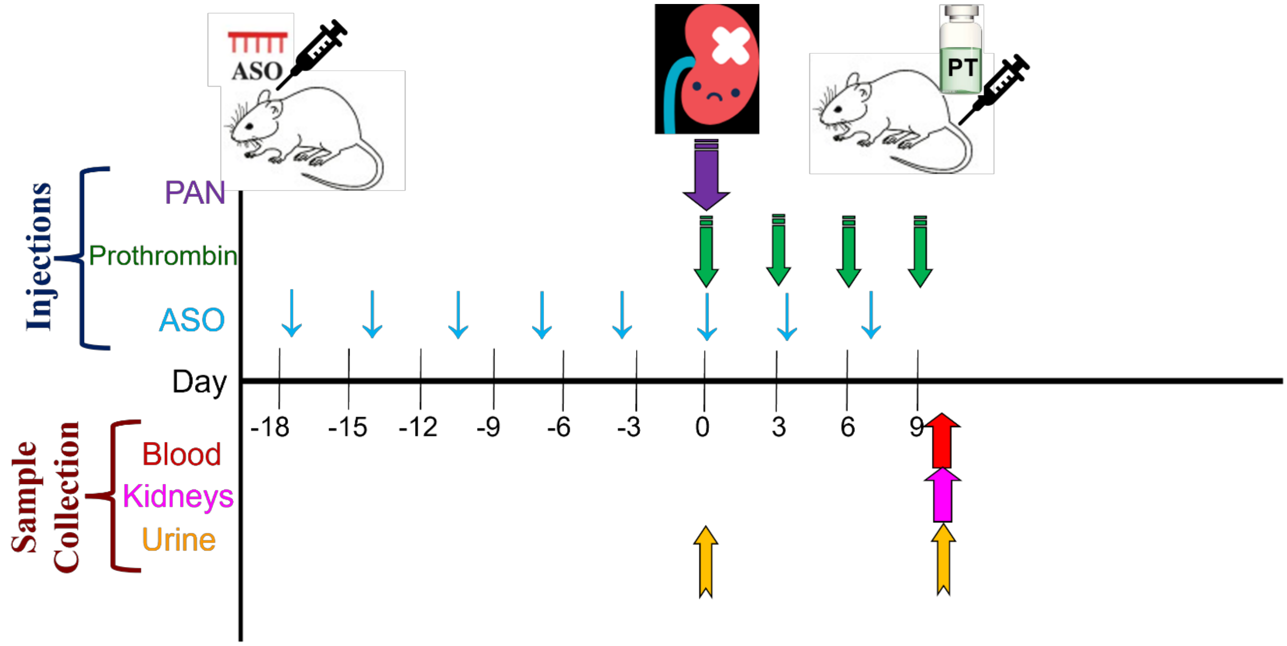


**Figure S1: Experimental Schema.** Puromycin Aminonucleoside (PAN) was delivered intravenously to induce nephrosis on day 0 (purple arrow), control rats received intravenous saline instead. Hyper-prothrombinemia (HiPT) was modeled using intravenously delivered human prothrombin on days 0, 3, 6, and 9 (green arrows). Hypo-prothrombinemia (LoPT) was modeled by twice weekly delivery of rat prothrombin-specific antisense oligonucleotide (ASO) 401027 beginning on day -17 (blue arrows). See methods for doses used. Urine was collected on days 0 and 10 (yellow arrows). On day 10, after urine collection, the rats were exsanguinated (red arrow) and their kidneys were collected (pink arrow).

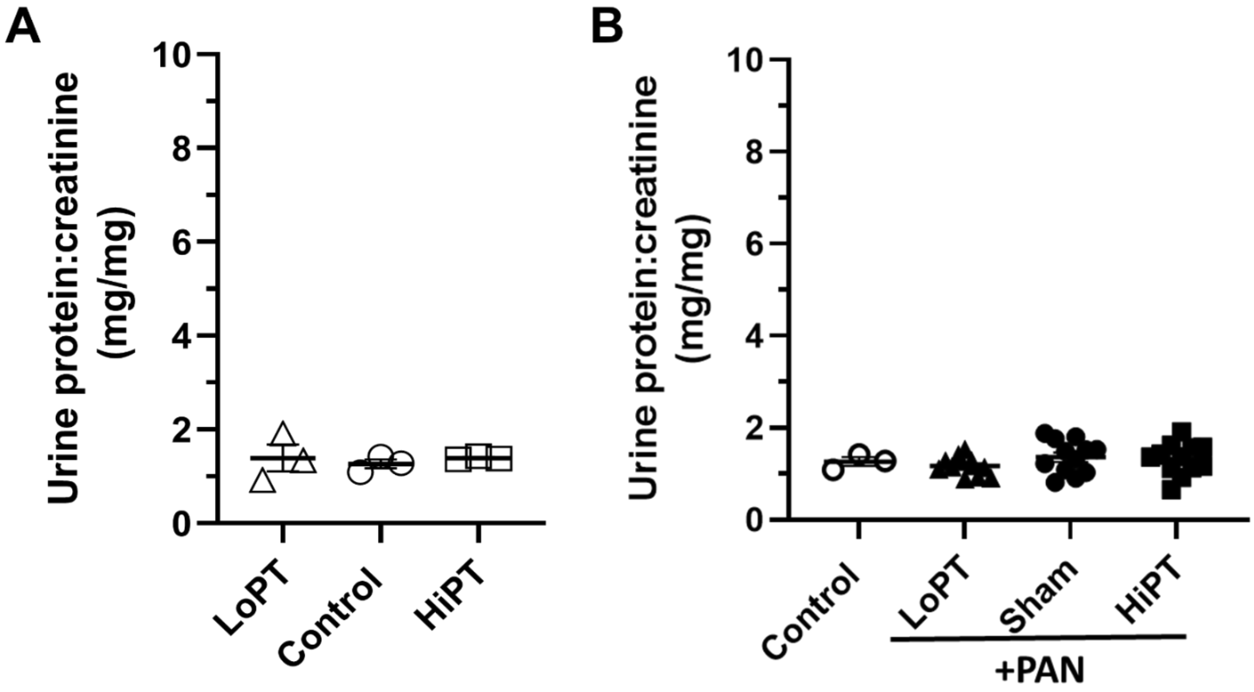


**Figure S2: Proteinuria was unchanged on Day 0.** Prothrombin modulated healthy rats (**A**) and PAN-NS rats (**B**) had equivalent day 0 urinary protein. *n*=3-12 per group

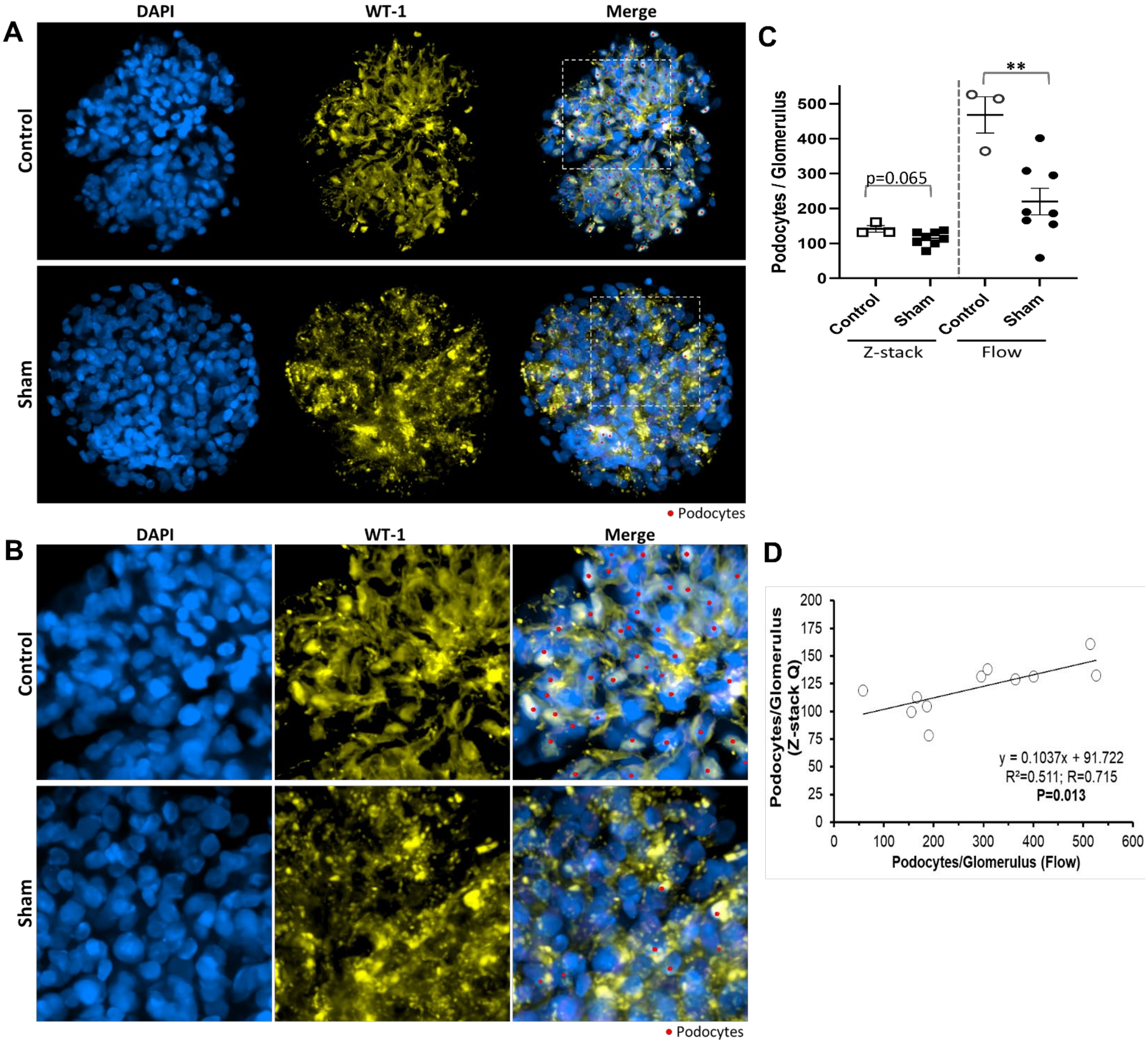


**Figure S3: Histologic Podocyte Counts Correlated with Flow Cytometry Podocyte Counts.** Isolated rat glomeruli were immunofluorescently labeled with DAPI (non-specific nucleus stain, blue) and anti-WT-1 antibody (specific to podocyte nuclei, yellow); z-stack images (1 µm z-step) were collected from each randomly selected glomerulus. (**A**) Representative images show 3D volume projections of cropped z-stacks showing only 6 consecutive z-slices for best visibility, a red marker is shown at the 3D center of each WT-1-positive nucleus; however, some partially visible WT-1-positive nuclei may not be apparent in this projection due to cropping in the z-dimension. (**B**) Magnification of inset area shown in panel A. (**C**) Podocyte counts were reduced in PAN-NS (Sham) vs. healthy control rats by both the histologic (each point represents averaged data taken from 15 glomeruli per rat) and flow cytometry (each point represents averaged data derived from ~2,500 glomeruli and 100,000 event counts per rat) methods but was only significant by flow cytometry. (**D**) There was a significant linear relationship between the histologic and flow cytometry methods. *n*=3-8 per group; $^{**}P<0.01$

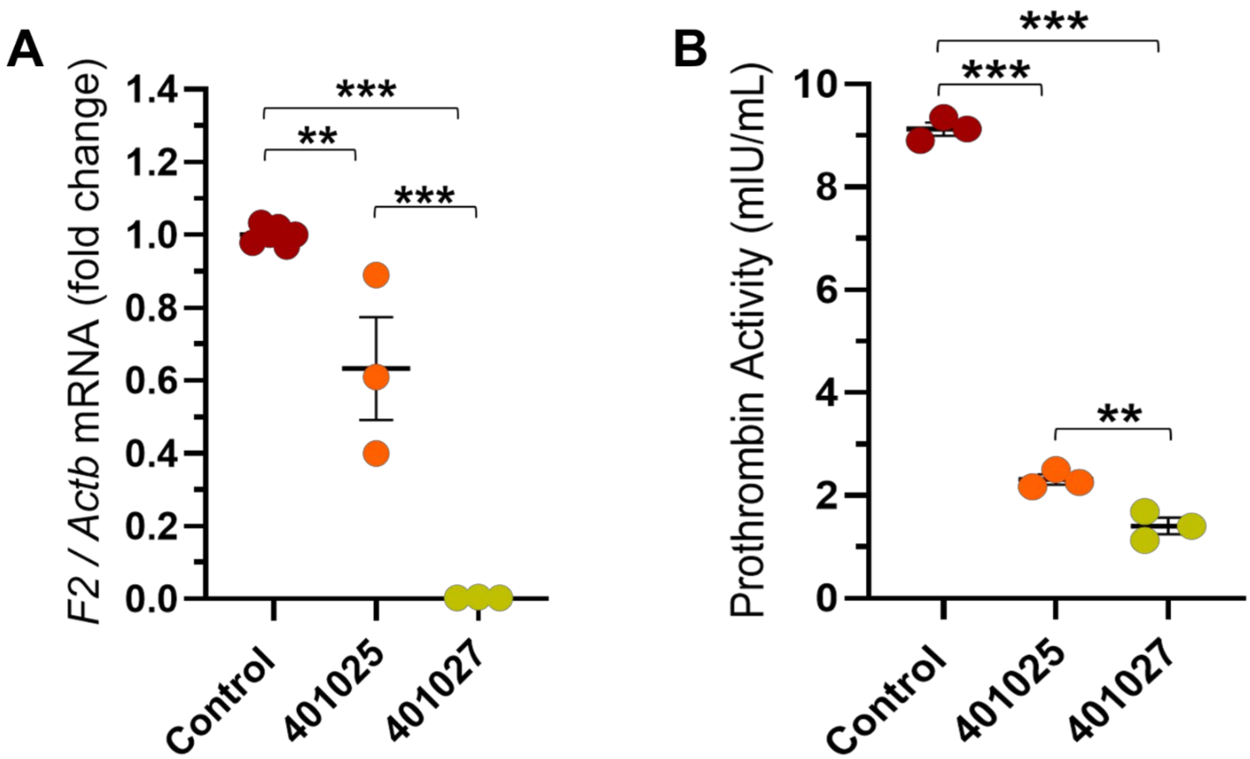


**Figure S4: Antisense Oligonucleotide 401027 Inhibited Rat *F2* Expression in Cultured Primary Rat Hepatocytes.** Primary rat hepatocytes were cultured in the presence of a control (scrambled) antisense oligonucleotide (ASO), ASO 401025 (75% complementary to rat *F2* 3' untranslated region (UTR)) or ASO 401027 (100% complementary to rat *F2* 3' UTR). (**A**) ASO 401027 essentially abolished *F2* expression (relative to β-actin (*Actb*)) whereas ASO 401025 less efficiently reduced *F2* expression (*n*=3-6 technical replicates per condition). (**B**) Both ASO 401025 and 401027 significantly reduced chromogenic (S-2238) prothrombin enzymatic activity in cultured primary rat hepatocyte protein lysates, but ASO 401027 was significantly better at reducing activity in comparison to ASO 401025 (*n*=3 technical replicates per condition). ***P*<0.01, ****P*<0.001

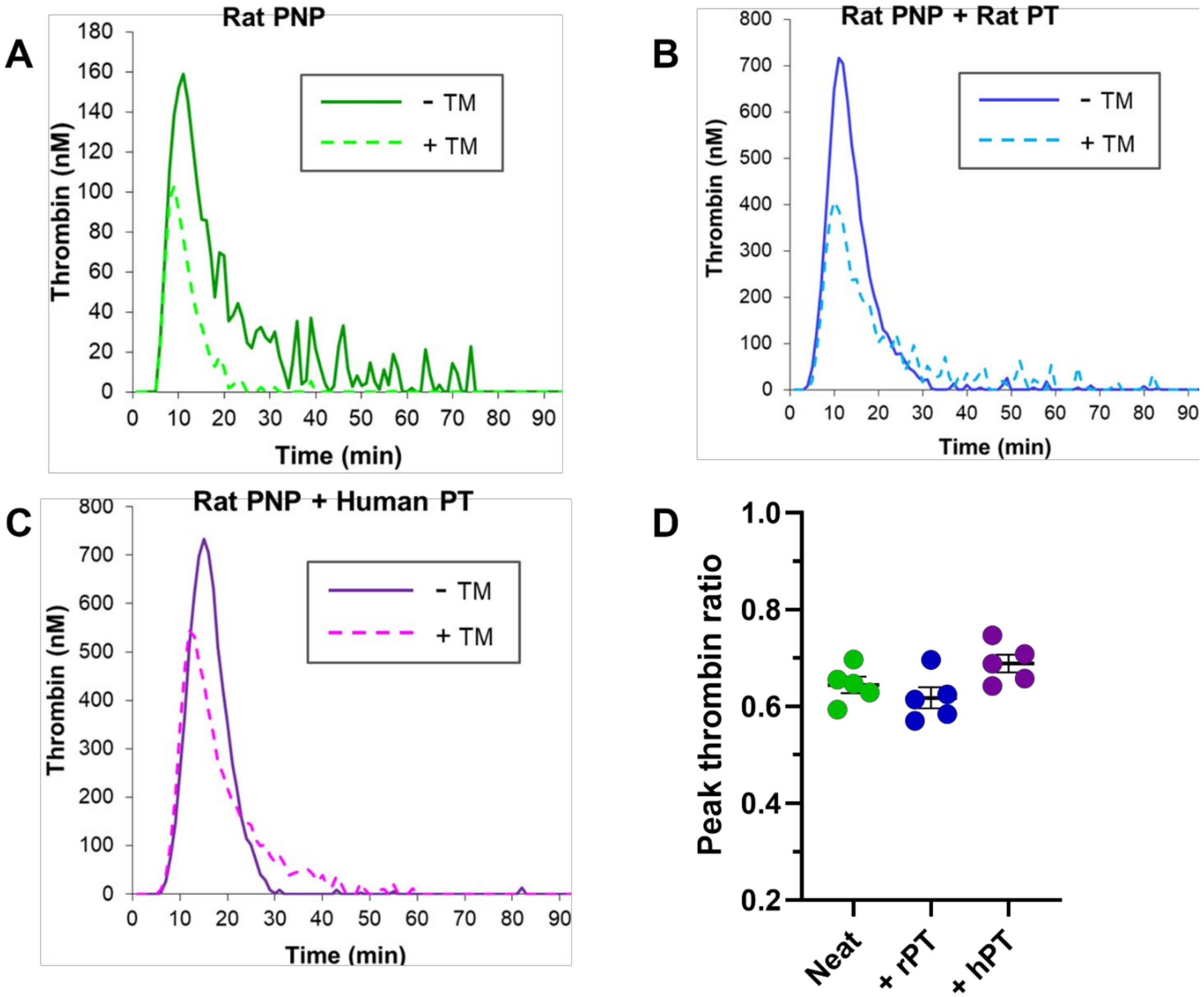


**Figure S5: Human Prothrombin Augmented Thrombin Generation in Rat Plasma and was Regulated by Rat Thrombomodulin.** Thrombin generation in rat pooled normal plasma (PNP, 3.5 μM prothrombin) was measured in the absence and presence of 200 nM rat thrombomodulin (TM). (**A**) Rat PNP spiked with prothrombin diluent. (**B**) Rat PNP was spiked to 7 μM prothrombin with rat prothrombin. (**C**) Rat PNP was spiked to 7 μM prothrombin with human prothrombin. (**D**) Peak thrombin ratio (+TM:-TM) from each condition demonstrates that rat TM effectively regulated both rat and human thrombin generation. *n*=5 technical replicates

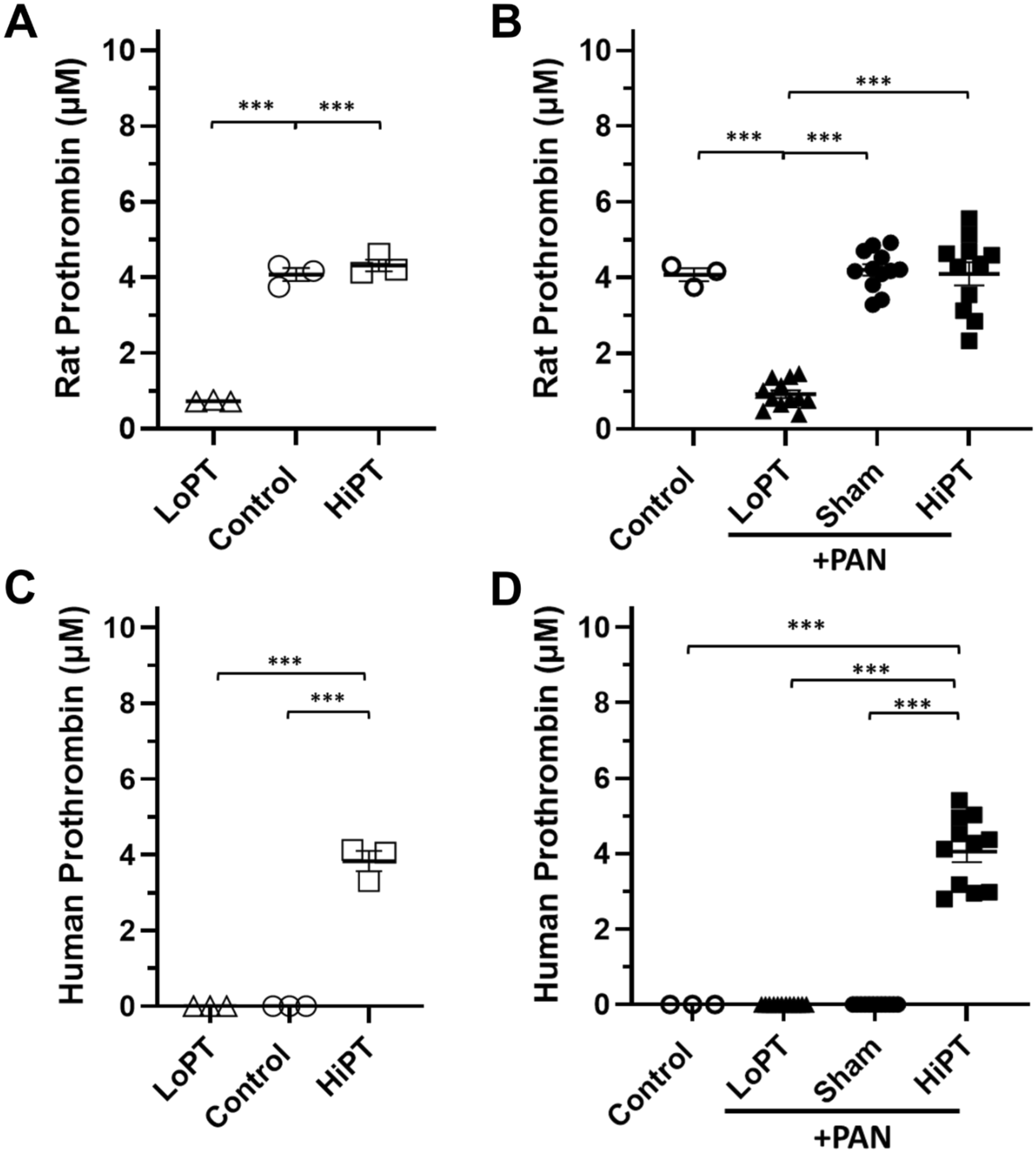


**Figure S6: Plasma Prothrombin Protein Quantification.** (**A**, **B**) Plasma prothrombin protein quantification in healthy control (**A**) and PAN-NS (**B**) rats using a rat-specific prothrombin ELISA. (**C**, **D**) Plasma prothrombin protein quantification in healthy control (**C**) and PAN-NS (**D**) rats using a human-specific prothrombin ELISA. For the HiPT groups, rat and human prothrombin quantities were added to arrive at total plasma prothrombin protein quantity as shown in Figure 3. $n$=3-12 per group; ***$P$<0.001

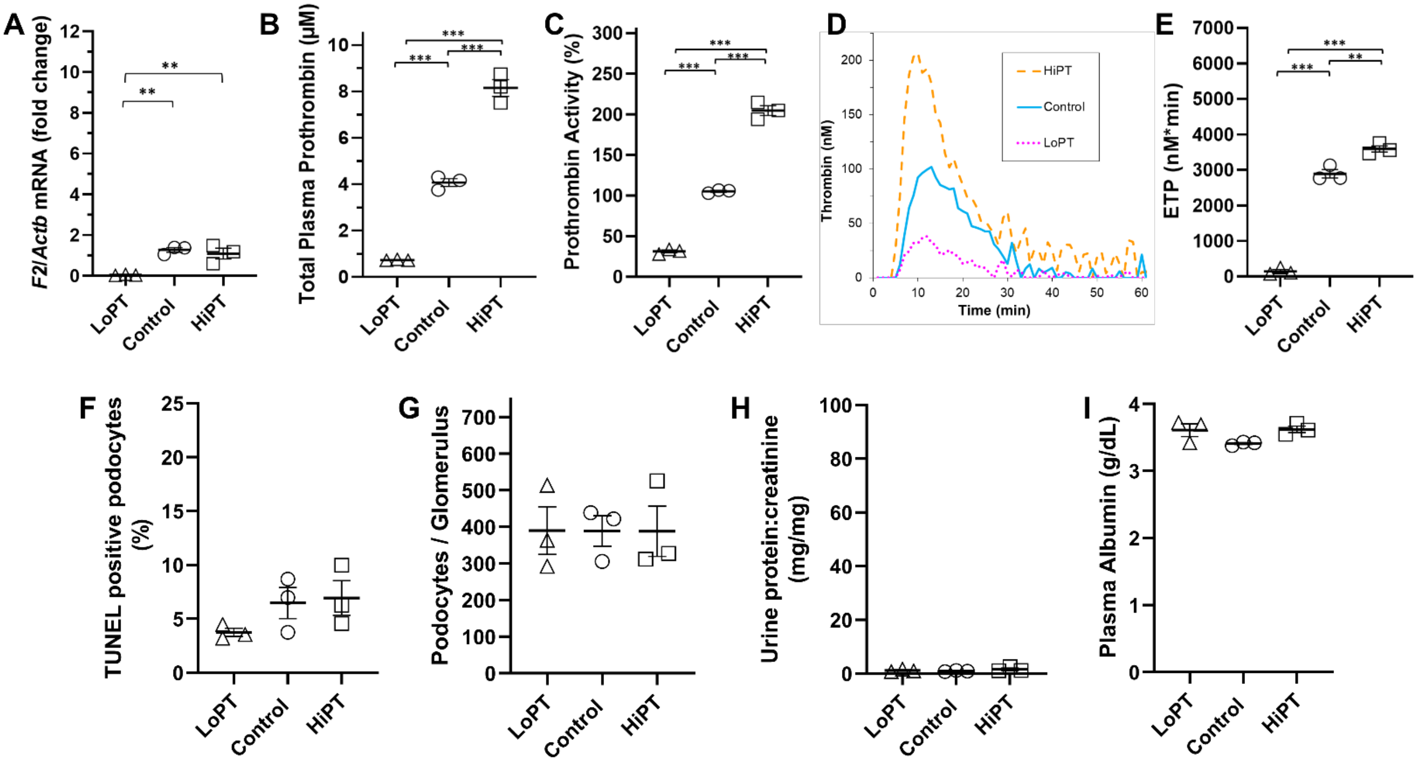


**Figure S7: Hypo- and Hyper-Prothrombinemia did not Alter Podocytopathy, Podocyte Counts, Proteinuria, or Plasma Albumin in Healthy Control Rats.** ASO 401027 and human prothrombin treatment identical to that administered to PAN-NS rats (Figure 3) significantly modulated hepatic *F2* expression (**A**) as well as plasma prothrombin protein quantity (**B**), enzymatic activity (**C**), and endogenous thrombin potential (ETP, **D**, **E**). However, these changes in circulating prothrombin levels did not significantly alter podocytopathy (**F**), podocyte counts (**G**), proteinuria (**H**), or plasma albumin (**I**). $n$=3 per group; **$P$<0.01, ***$P$<0.001

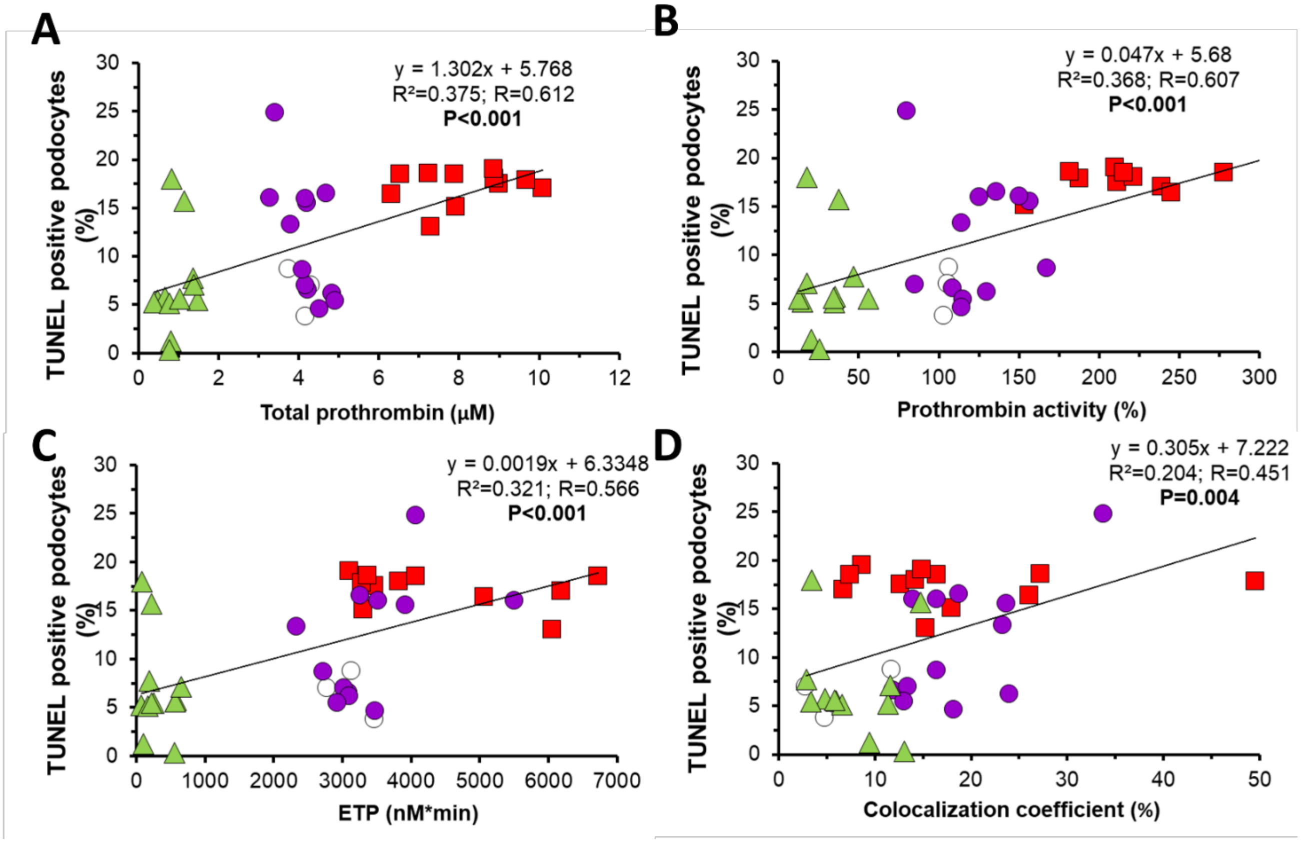


**Figure S8: Podocyte Injury Correlated with Prothrombin Levels and Thrombin-Podocyte Interactions.** Terminally injured, TUNEL-positive podocytes were significantly correlated with plasma prothrombin protein quantity (**A**), enzymatic activity (**B**), endogenous thrombin potential (ETP, **C**), and thrombin colocalization to podocytes (**D**). *n*=3-12 per group; ○: Control; ▲: LoPT (ASO-mediated hypo-prothrombinemia); ●: Sham; ■: HiPT (prothrombin infusion-mediated hyper-prothrombinemia)

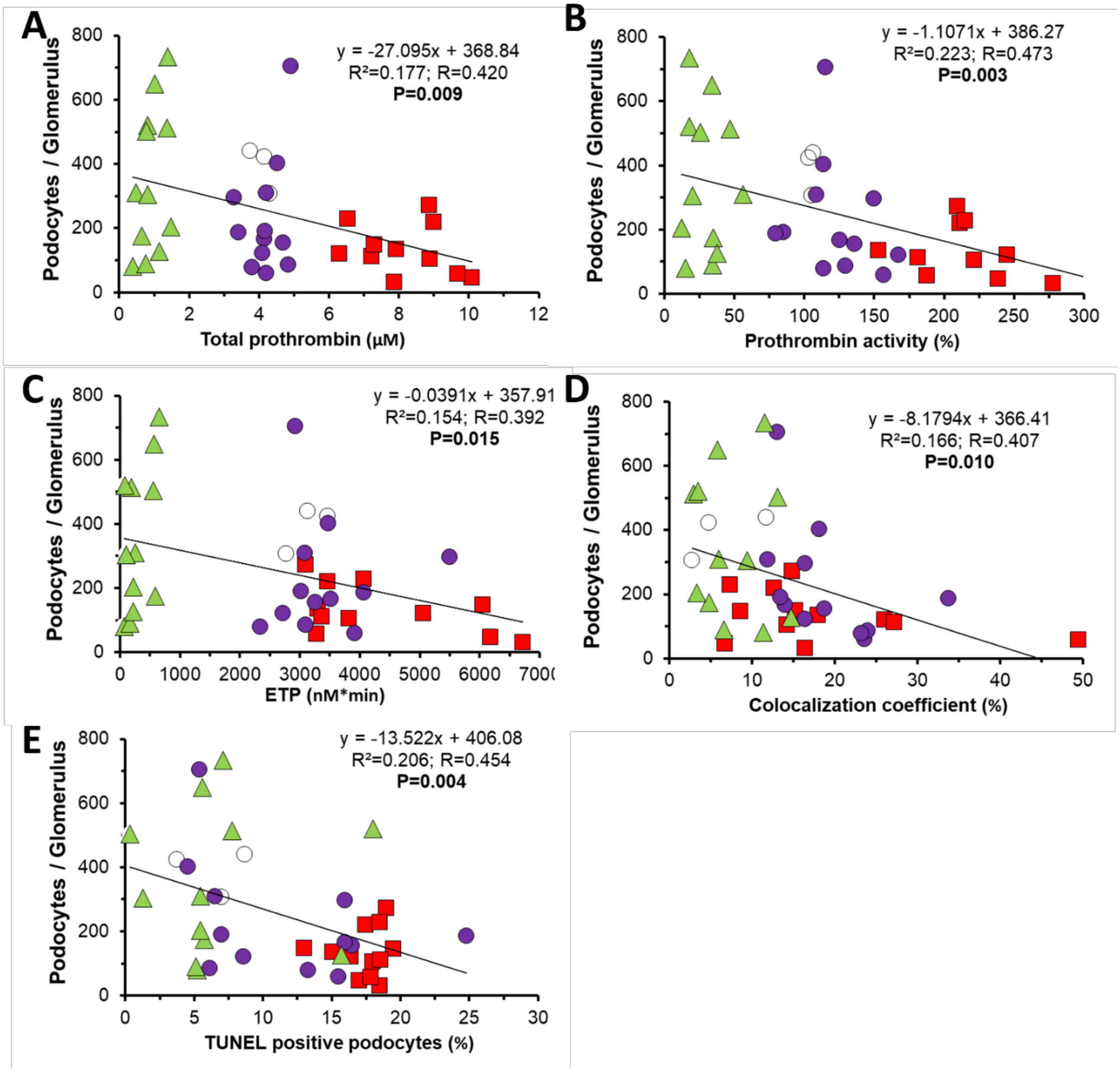


**Figure S9: Podocyte Survival Correlated with Prothrombin Levels, Thrombin-Podocyte Interactions, and Podocyte Injury.** Podocytes per glomerulus was significantly correlated with plasma prothrombin protein quantity (**A**) and enzymatic activity (**B**), but not endogenous thrombin potential (ETP, **C**). Podocyte counts were also significantly correlated with thrombin colocalization to podocytes (**D**) and TUNEL-positive podocytes (**E**). *n*=3-12 per group; ○: Control; ▲: LoPT (ASO-mediated hypo-prothrombinemia); ●: Sham; ■: HiPT (prothrombin infusion-mediated hyper-prothrombinemia)

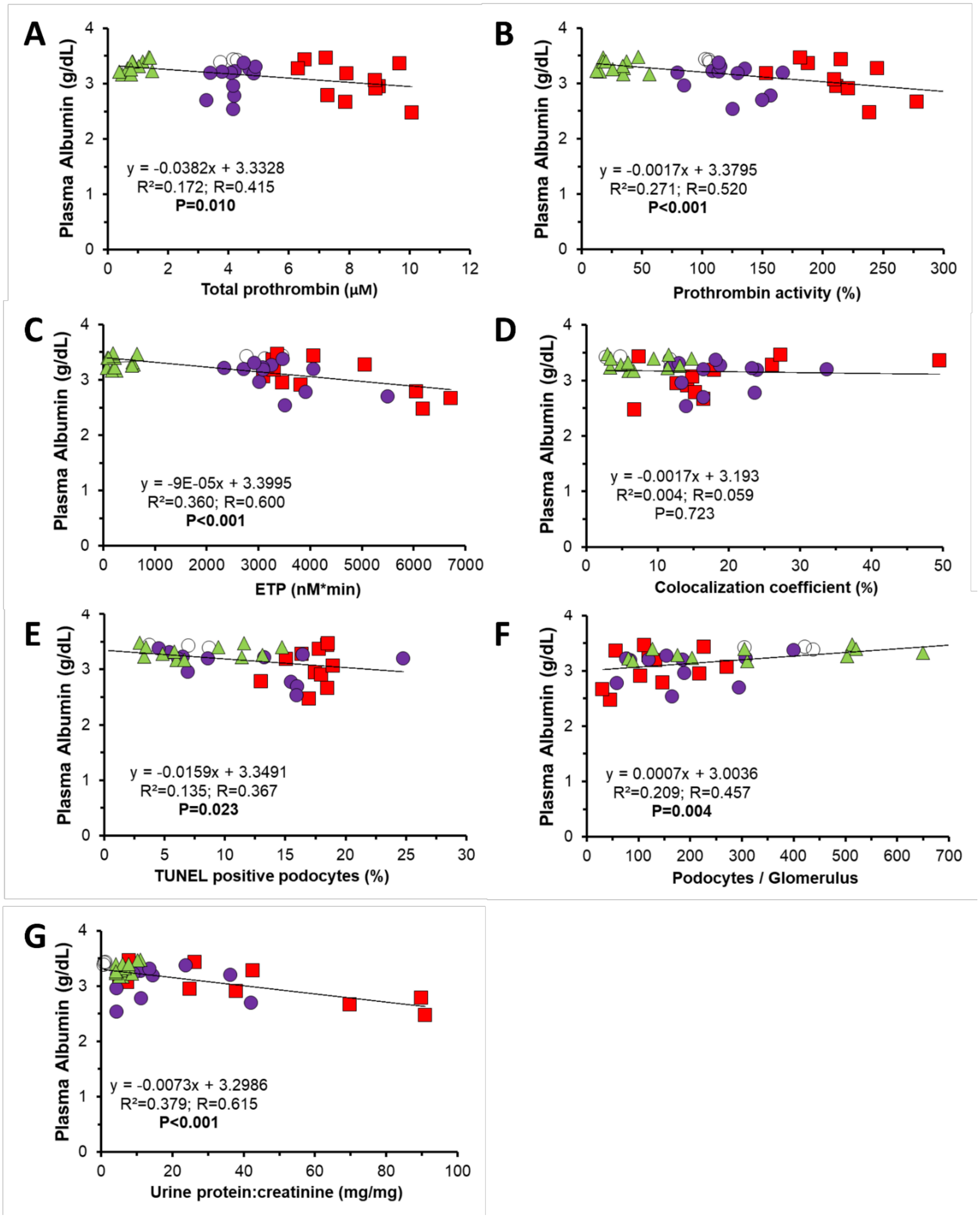


**Figure S10: Plasma Albumin Correlated with Prothrombin Levels, Podocyte Injury, Podocyte Survival, and Proteinuria.** Plasma albumin was significantly correlated with plasma prothrombin protein quantity (**A**), enzymatic activity (**B**), and endogenous thrombin potential (ETP, **C**). Thrombin colocalization to podocytes (**D**) was not related to plasma albumin level. TUNEL-positive podocytes (**E**), podocytes per glomerulus (**F**), and proteinuria (**G**) were significantly correlated with plasma albumin. *n*=3-12 per group; ○: Control; ▲: LoPT (ASO-mediated hypo-prothrombinemia); ●: Sham; ■: HiPT (prothrombin infusion-mediated hyper-prothrombinemia)

### SUPPLEMENTAL MOVIES

**Video 1: 2D Scan through a Representative 3D Reconstructed Control Glomerulus.** Isolated glomerulus immunofluorescently labeled with DAPI (non-specific nucleus stain, blue) and WT-1 antibody (specific to podocyte nuclei, yellow). Z-stack images (1 μm step) were collected, 3D reconstructed, and a red marker was placed at the 3D center of each WT-1-positive nucleus. The video scans through the 3D z-stack with each frame representing a 1 μm thick section.

**Video 2: 2D Scan through a Representative 3D Reconstructed Sham Glomerulus.** Isolated glomerulus immunofluorescently labeled with DAPI (non-specific nucleus stain, blue) and WT-1 antibody (specific to podocyte nuclei, yellow). Z-stack images (1 μm step) were collected, 3D reconstructed, and a red marker was placed at the 3D center of each WT-1-positive nucleus. The video scans through the 3D z-stack with each frame representing a 1 μm thick section.

**Video 3: Rotational View of a Representative 3D Reconstructed Control Glomerulus.** Isolated glomerulus (same as in Movie M1) immunofluorescently labeled with DAPI (non-specific nucleus stain, blue) and WT-1 antibody (specific to podocyte nuclei, yellow). Z-stack images (1 μm step) were collected, 3D reconstructed, and a red marker was placed at the 3D center of each WT-1-positive nucleus. The video rotates the 3D reconstructed glomerulus around the x-, then y-axis. Flattening of the glomerulus in the z-dimension is an artifact of the coverslip used to prepare the glomeruli for imaging.

**Video 4: Rotational View of a Representative 3D Reconstructed Sham Glomerulus.** Isolated glomerulus (same as in Movie M2) immunofluorescently labeled with DAPI (non-specific nucleus stain, blue) and WT-1 antibody (specific to podocyte nuclei, yellow). Z-stack images (1 μm step) were collected, 3D reconstructed, and a red marker was placed at the 3D center of each WT-1-positive nucleus. The video rotates the 3D reconstructed glomerulus around the x-, then y-axis. Flattening of the glomerulus in the z-dimension is an artifact of the coverslip used to prepare the glomeruli for imaging.
